# Alignment between Duplex Sequencing and transgenic rodent mutation assay data in the assessment of in vivo NDMA-induced mutagenesis

**DOI:** 10.1101/2025.04.29.651210

**Authors:** Anne L. Ashford, Daniela Nachmanson, John W. Wills, Jacob E. Higgins, Thomas H. Smith, Kevin C. Vavra, Farah Dahalan, Jonathan Howe, Joanne M. Elloway, Jesse J. Salk, Ann Doherty, Anthony M. Lynch

## Abstract

The nitrosamine N-nitrosodimethylamine (NDMA) is a mutagen and rodent carcinogen that has been identified as a process impurity in some commercially available medicines, leading to market withdrawals and new impurity control measures. Error-corrected DNA sequencing techniques, such as Duplex Sequencing (DS), have error rates low enough to revolutionise genetic toxicology testing by directly measuring *in vivo* mutagenesis within days of exposure. Here, DS was performed on liver samples from an OECD-compliant, Transgenic Rodent Gene Mutation Assay (TGR) conducted under GLP standards. Muta™Mouse specimens were orally dosed with NDMA using either a repeat-dose 28-day regimen (0.02–4 mg/kg(bw)/day) or single bolus doses of either 5 or 10 mg/kg(bw) administered on day one. Dose-dependent increases in mutation frequency were detected by DS in liver, enabling a No-Observed Genotoxic Effect Level (NOGEL) of 0.07 mg/kg(bw)/day to be determined, supported by mechanistic analyses of trinucleotide mutation spectra. Benchmark dose (BMD) modelling determined similar BMD_50_ values from either DS or TGR, demonstrating concordance across the two techniques albeit with greater precision from DS due to smaller inter-animal variation. DS offers a fundamental change in mutagenicity assessments enabling more precise point-of-departure determinations with mechanistic clarity and 3Rs advantages compared to the standard TGR approach.

## INTRODUCTION

N-Nitrosamines, including N-Nitrosodimethylamine (NDMA), are a class of compounds that are known for being potential genotoxic carcinogens. N-nitrosamines can be detected in the environment, e.g. in water and certain foods, and can be formed as process impurities in some industrial and consumer products (ATSDR, FDA guidance). NDMA is a small, potent genotoxic nitrosamine, that can cause DNA damage by forming methyl adducts with DNA bases (DNA alkylation), primarily at the O6-position of guanine which can lead to miscoding during DNA replication, potentially resulting in C:G -> T:A transition mutations (Delker et al. 2008; Loechler et al. 1984; Shane et al. 2000; Souliotis et al. 1998; Wang et al. 1998). NDMA is a rodent carcinogen and IARC, ATSDR and WHO have classified NDMA as a Group 2A carcinogen, indicating that it is probably carcinogenic to humans (IARC 1978).

Recently, N-nitrosamine impurities have been detected in some commercial medicines. This has led to the recall of some pharmaceutical products worldwide and increased regulatory scrutiny regarding the assessment of trace nitrosamine impurities in commercial medicines (Nudelman 2023). Potential nitrosamine impurities are a diverse chemical group, including small nitrosamines, such as NDMA, but also novel, larger drug-like nitrosamines (Nitrosamine Drug Substance-Related Impurities, NDSRIs). Nitrosamine impurities can be present in raw materials or be formed during manufacture or under certain storage conditions (Moser et al. 2023; Schlingemann et al. 2022). A review of the mechanisms by which certain medicines were contaminated with NDMA impurities can be found in Schlingemann et al. 2022.

The *in vivo* mutagenic potential of compounds, including N-nitrosamines, can be determined using Transgenic Rodent Somatic and Germ Cell Gene Mutation Assays (TGR mutation assays). In this assay, commercially available transgenic rat or mouse strains (e.g., Big Blue® rodents or Muta™mouse) contain multiple copies of chromosomally integrated phage shuttle vectors which are used to detect chemicals that may induce gene mutations in somatic and germ cells (OECD Test No. 488, 2022). These validated *in vivo* mutagenicity assays are considered appropriate to qualify the mutagenic potential of impurities for regulatory purposes (ICH M7(R2) 2023). The TGR mutation assays are highly concordant with the 2-year rodent bioassay for genotoxic carcinogens (Lambert et al. 2005; Zeller et al. 2018). The basis for the TGR mutation assay is predicated on last century technology and relies on indirect measurements of phenotypic mutants using prokaryotic reporter genes: The mutant gene frequency is based on phenotypic changes induced by phage transduction in the specific bacterial cell strains, and formation of plaques in the absence or presence of selection. In addition, TGR studies are labour intensive with limited study slot availability. *In vitro* mutagenic potential can be determined using the Ames test but the ability of the Ames test to provide a sensitive method to detect mutagenicity of nitrosamines has been under investigation (Heflich et al. 2024; Li et al. 2023; Tennant et al. 2023; Trejo-Martin et al. 2022). Major health authorities have issued guidance for Enhanced Ames Test (EAT) conditions to improve sensitivity for the detection of mutagenic nitrosamine drug substance-related impurities (NDSRIs), but, at the time of writing, a negative result does not allow control of the NDSRI as a non-mutagenic impurity (EMA 2020). A more informative and biologically relevant approach could be to detect mutations using DNA sequencing of the mammalian genome which would provide not only a direct measure of mutations but also additional information on mutation type and sequence context.

Next Generation Sequencing (NGS) has revolutionised the speed and throughput of DNA and genome sequencing. Short read sequencing has an error rate of 0.1-1% per base which is acceptable for most applications but raises problems for genetic toxicology assays, where whole cell populations are analysed in bulk for mutation burden. In these assays, unique mutations found in a single cell at a single genomic location and among many millions of unmutated bases, are obscured by the inherent error rate associated with sequencing. Therefore, ultra-sensitive methods with very low error rates are required. Such techniques are termed error-corrected NGS (ecNGS) and use a variety of methods to reduce the error rate by 10,000-fold or more (Marchetti et al. 2023; Salk and Kennedy 2020; Salk et al. 2018). Some of the most accurate methods to-date are based on Duplex Sequencing (DS) (Kennedy et al. 2014; Schmitt et al. 2012), which tag double stranded DNA with DNA barcodes, allowing sequencing reads to be traced back to a single starting DNA molecule, including its forward or reverse strand. This enables the creation of single and double strand consensus sequences. Since the probability of an error occurring at the exact same location on both strands is tiny, double strand consensus sequences effectively eliminate nearly all sequencing errors, reducing the error rate down to an estimated 1x10^-8^ – 1x10^-7^, and are sufficiently sensitive to detect the ultra-rare mutation events required by genetic toxicology assays for mutagenesis (Valentine et al. 2020). Indeed, DS has been utilised to assess the *in vivo* mutagenic potential of known mutagens, e.g. Benzo[a]pyrene, Benzo[b]fluoranthene, N-ethyl-N-nitrosourea, procarbazine (Dodge et al. 2023; LeBlanc et al. 2022; Schuster et al. 2024; Smith-Roe et al. 2023; Valentine et al. 2020), and the nitrosamine, N-Nitrosodiethylamine (NDEA) (Bercu et al. 2023; Zhang et al. 2024). To assess the concordance between TGR and DS, an important assessment is benchmark dose (BMD) analysis on TGR and DS data that have both been generated from the bacterial transgene as this is the most like-for-like comparison: Whilst some of these studies compared TGR and DS dose-responses using BMD modelling, none of these analyses used ‘matched’ DS data from the TGR bacterial transgene itself.

In this study, we used DS to further characterise NDMA-induced mutagenesis *in vivo* over a broad range of doses using tissue samples from an OECD 488 compliant TGR mutation assay (Lynch et al. 2024). The original TGR study reported the analysis of mutant frequency responses in multiple tissues (liver, lung, kidney, spleen, stomach, bladder, and bone marrow) and included additional genotoxicity measurements i.e., the micronucleus test (OECD Test guideline 474) and Pig-a assay (OECD Test guideline 470) along with liver tissue pathology and toxicokinetics (Lynch et al. 2024). The study included a 28-day, daily dosing regimen alongside two single-dose treatments, the latter providing doses equivalent to the cumulative doses of two of the repeat-dose treatments. The original TGR mutation assay provided an indirect measure of mutation burden via the detection of *lacZ* mutant gene frequency. The purpose of the current study was to perform DS analysis on matched liver samples taken from this study. DS data was generated using either a ‘mouse mutagenesis panel’, which directly measured mutations contained in the DNA from representative regions established across the mouse genome, or a ‘*lacZ* mutation panel’ that specifically targeted the TGR transgene. This enabled the data from the two panels and the original TGR *lacZ* mutant frequency responses to be compared. Liver tissue was chosen as potent nitrosamines, including NDMA, require metabolism via cytochrome p450 2E1 enzymes, to generate the DNA-reactive proximate mutagen (Godoy et al. 1978; Yamazaki et al. 1992). As such liver was the target organ with the highest mutation burden in existing transgenic rodent studies (Gollapudi et al. 1998; Jiao et al. 1997; Lynch et al. 2024; Shane et al. 2000; Suzuki et al. 1996) and it is the most susceptible organ to NDMA-induced carcinogenesis in rodent (Peto et al. 1991a; Peto et al. 1991b). The aim of the study was to characterise DS for the assessment of mutagenesis, robustly bridge to other existing regulatory assays using BMD modelling as a comparator and support the development and integration of ecNGS approaches into *in vivo* test guidelines for regulatory purposes.

## METHODS

### Isolation of genomic DNA

Genomic DNA was isolated from approximately 25mg of frozen liver tissue using a DNeasy Blood and Tissue Kit (Qiagen) following the manufacturer’s protocol. The tissue was homogenised using the FastPrep-24 bead homogneisation system with Matrix D tubes (MP Biomedicals) in Buffer ATL (Qiagen). The proteinase K incubation was performed at 37°C for 2 hours and the optional RNase A step was included. DNA concentration and quality was determined using Nanodrop (ThermoFisher), Qubit high sensitivity DNA assay kit (Thermo Fisher) and Tapestation Genomic DNA ScreenTape Analysis (Agilent).

### Duplex Sequencing library preparation

For each sample, 760ng of genomic DNA was split to generate two Duplex Sequencing libraries; 160ng to sequence the *lacZ* multicopy transgene and 600ng to sequence using the Mouse Mutagenesis Panel (TwinStrand). This was to prevent over-sequencing of the *lacZ* gene due to the multi-copy nature of the transgene in the MutaMouse model. Duplex Sequencing libraries were prepared as described before (Valentine et al. 2020), with the exception that genomic DNA was enzymatically fragmented instead of sonicated. Briefly, enzymatically fragmented DNA was end-repaired and A-tailed, followed by ligation of DuplexSeq™ adapters containing unique molecular identifiers (UMI). Library conditioning was performed using a mixture of glycosylases to remove damaged DNA prior to amplification. Following indexing PCR, target regions of DNA were enriched by hybrid capture using either the *lacZ* or Mouse Mutagenesis Panel.

The *lacZ* custom panel targets the entire coding sequence of the *lacZ* gene, this sequence was derived from a *de novo* MutaMouse assembly of the λgt10 genome (Meier et al. 2019). Meanwhile, the Mouse Mutagenesis Panel targets 20 endogenous mouse genomic loci, each 2.4 kb in size, spread across all autosomes. Nine of the targets are located within genic regions and 11 within intergenic regions; not all genic targets are contained within the coding sequence of the gene and not all genes will be expressed in all tissues, see additional details as described before (Dodge et al. 2023). Following hybridization selection, libraries were purified with streptavidin magnetic beads. After washes, additional PCR was performed, followed by another round of hybridization, capture, washes, and a final round of PCR. Final libraries were quantified with a Qubit fluorometer and dsDNA HS reagents, and fragment size was assessed using an Agilent TapeStation system and high sensitivity D1000 reagents prior to pooling. Library pools were sequenced using paired-end 150bp sequencing on an Illumina NovaSeq 6000

### Duplex Sequencing bioinformatic processing

Bioinformatic processing was carried out as per the methods described in Valentine et al (Valentine et al. 2020). Alignment was performed using the BWA aligner version v0.7.17 (Li 2013). For the Mouse Mutagenesis panel the alignment was against the *Mus musculus* reference genome mm10 (GCA_000001305.2), while for the *lacZ* panel, the alignment was against a concatenation of *mm10* and MutaMouse λgt10 reference genome (Meier et al. 2019). Post-processing of the duplex consensus alignments included balanced overlap hard clipping to limit the generation of double-counted variants during the subsequent variant calling process. Duplex consensus alignments from the Mouse Mutagenesis Panel were filtered to retain only those that were unambiguously from the *Mus Musculus* genome assembly using the taxonomic classifier Kraken v2.1.2 (Wood et al. 2019). Somatic and germline variants were called using VarDictJava in tumour-only mode v1.8.3 (Lai et al. 2016).

### Somatic mutation filtering

To ensure that only variants resulting from mutagenesis were considered, several filtering steps were implemented. For the Mouse Mutagenesis panel, we applied a germline threshold with a variant allele frequency (VAF) cutoff of 1%. However, for the *lacZ* gene, due to its multicopy nature (∼40–50 copies), we used a lower VAF threshold of 0.1%. Identical mutations appearing in more than one molecule within the same sample were considered to be derived from clonal expansion and were counted only once. Manual visual inspection was performed for clonal variants that did not present the expected germline VAF and for those that were found in three or more unique samples. This process led to the removal of 36 unique variants across all Mouse Mutagenesis Panel samples and 3 from the *lacZ* panel samples. Nearly all of these recurrent variants were attributed to misalignment in a single region on chromosome 1, The remaining variants were due to long structural variants or insertions/deletions that partly overlapped the targeted regions. The latter recurrent variant filtering steps represented removal of 2.5% of variants in the Mutagenesis panel and 0.4% of variants in the *lacZ* panel.

### Statistical testing

For the DS mutation data, pairwise comparisons of mutation frequency were performed using the glm function in R (version 3.6.1). To assess the statistical significance of differences between groups, a generalized linear model (GLM) was constructed with a quasi-Poisson error distribution, which accounts for overdispersion in the mutation count data. The natural logarithm of the number of informative duplex bases was included as an offset in the model. The p-values derived from the GLM were adjusted for multiple hypothesis testing using the Benjamini-Hochberg procedure to control the false discovery rate.

For comparison of the in-group variability (Fig 2c-d), statistical comparisons were performed using a paired Wilcoxon test, and p-values were adjusted for multiple hypothesis testing using the Benjamini-Hochberg procedure.

**Fig. 1.**
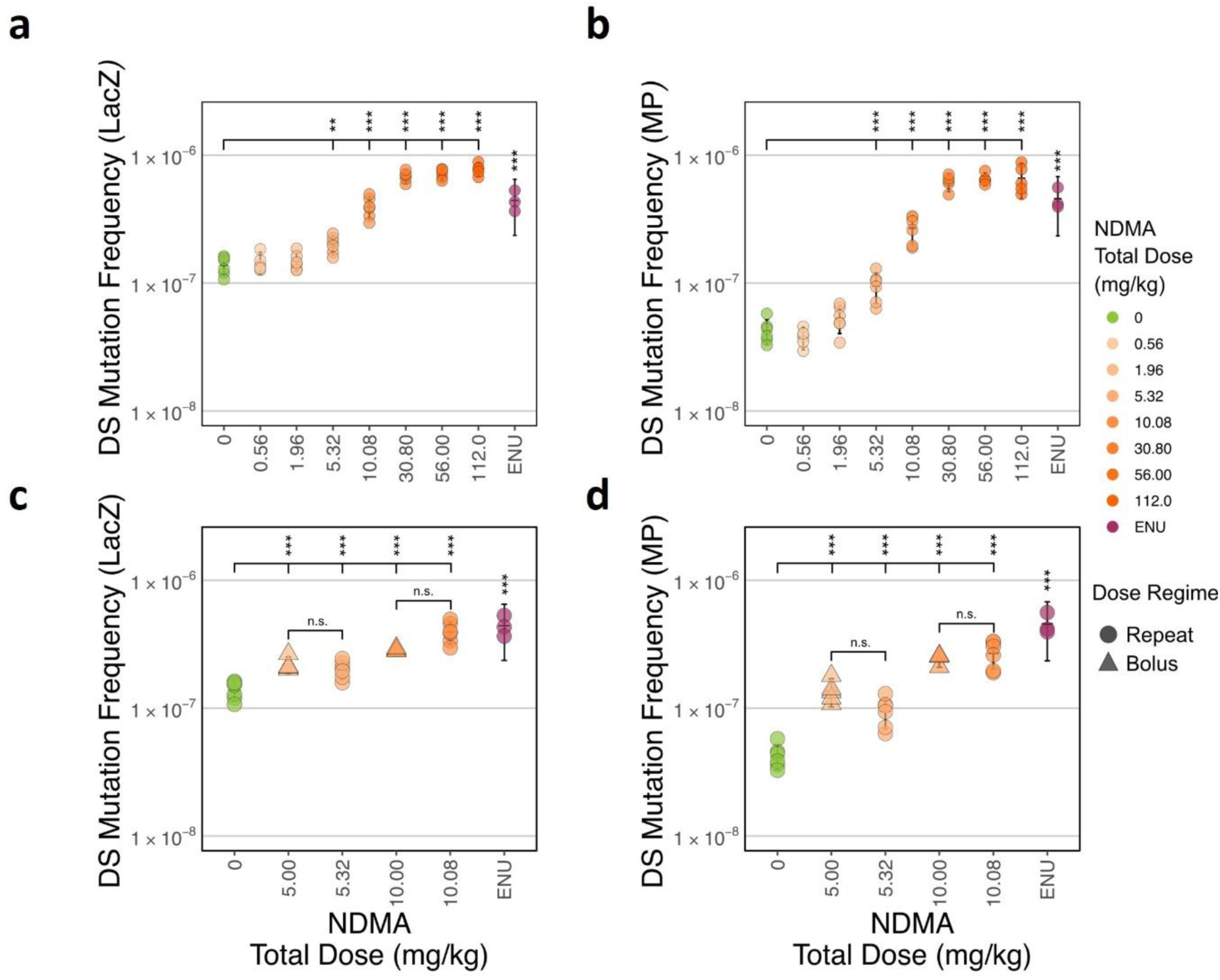
Dose-dependent mutation frequency induction in the Muta™Mouse liver following NDMA exposure using Duplex Sequencing. Panels **a** and **c** display mutation frequencies (MF) from Duplex Sequencing (DS) of the LacZ Panel, while panels **b** and **d** show results from the Mouse Mutagenesis (MP) panel. A and B illustrate repeat dosing only, while C and D include both repeat and bolus dosing. MF are shown across varying NDMA doses, with ENU as a positive control. Statistical significance is denoted by *** (p < 0.001), ** (p < 0.01), * (p < 0.05), and n.s. (not significant), with error bars representing 95% confidence intervals calculated using a t-distribution

**Fig. 2.**
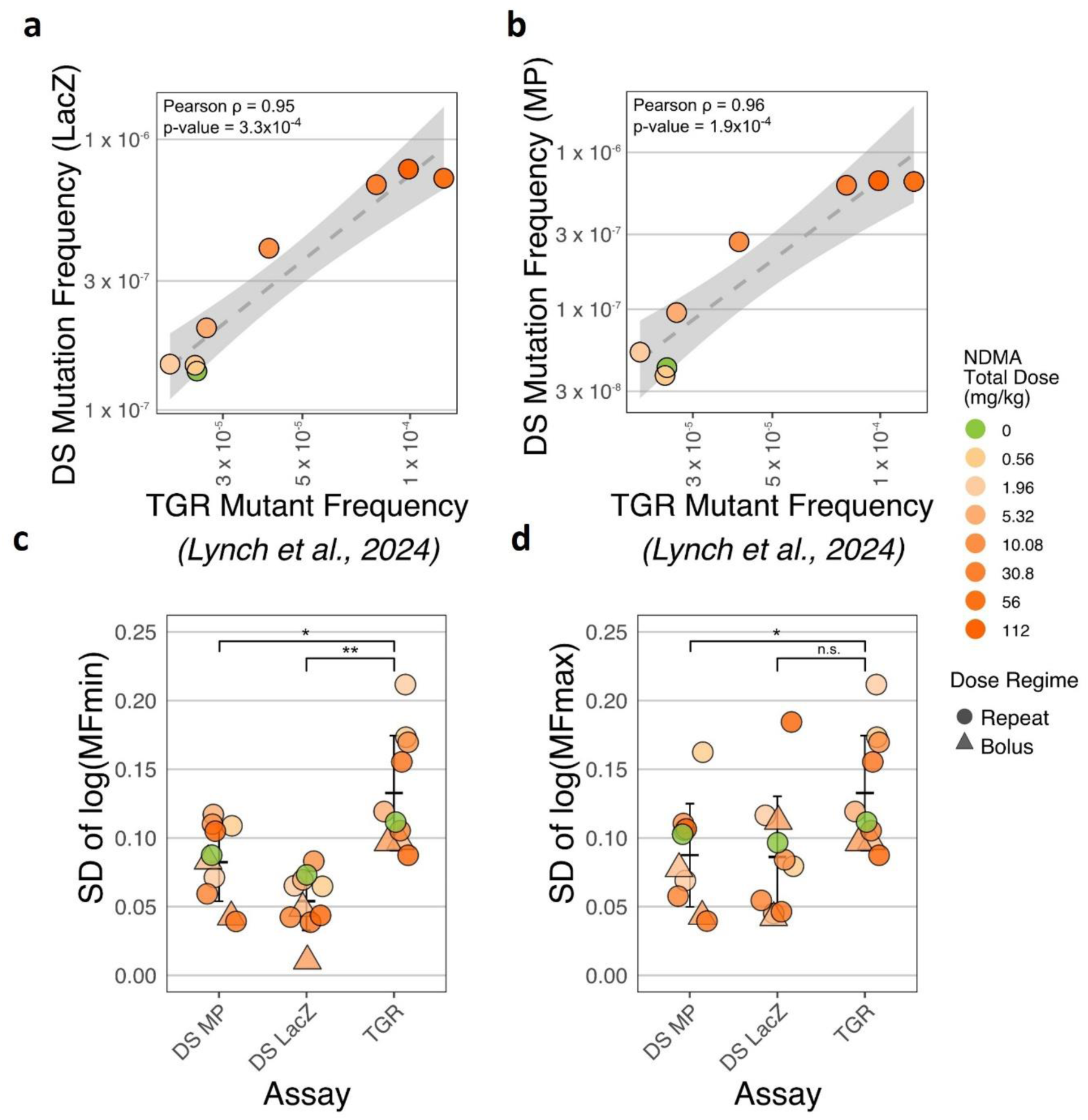
Comparison of TGR Muta™Mouse and Duplex Sequencing of the same liver tissues. Panels A and B show the correlation between TGR mutant frequency and Duplex Sequencing (DS) mutation frequency for **a** the *lacZ* panel and **b** mouse mutagenesis panel (MP). TGR measures the proportion of *lacZ* genes that harbour a functional mutation whereas Duplex Sequencing directly measures the frequency of all mutations (see text). The shaded grey areas represent the 95% confidence intervals. Pearson correlation coefficients (ρ) and p-values are indicated. Panels **c** and **d** display the in-group variability (standard deviation) of log-transformed mutation frequencies for DS and TGR assays. Panel **c** uses the “min” method to correct for clonality, where identical mutations at the same position are counted only once, while Panel **d** uses the “max” method, counting multiple identical mutations at the same position (potential clones) as independent mutation events. Statistical significance is denoted by * (p < 0.05) and ** (p < 0.01), with error bars representing 95% confidence intervals calculated using a t-distribution.

### Generation of predicted mutation data for *lacZ* panel

For the correction for trinucleotide context content to predict mutation frequency and substitution types, to determine whether differences in trinucleotide context abundance between the panels could explain the observed mutation frequency (MF) differences, we computed a context-corrected MF for the Mutagenesis panel. Specifically, the data were first separated by trinucleotide context, and then the mutation depth and informative duplex base counts were scaled using a context-specific ratio (LacZ:Mouse Mutagenesis panel context ratio) derived from the trinucleotide context counts. The corrected overall MF, or simple spectra MF, was calculated using these adjusted values (see Code for further details).

### Trinucleotide spectra analysis

Trinucleotide spectra were computed by first pooling all the mutations from animals within the same treatment group. For each unique trinucleotide context, we calculated the mutation frequency by dividing the number of observed mutations in a specific trinucleotide context by the total number of times that particular reference context was sequenced. For example, the mutation frequency for C[C>G]T mutations was computed by dividing the number of C[C>G]T mutations by the total number of CCT contexts sequenced. Subsequently, we calculated the proportion of each trinucleotide context mutation by dividing its mutation frequency by the sum of all mutation frequencies. This normalization step accounts for variability in sequencing depth across different trinucleotide contexts.

To perform statistical comparisons of the trinucleotide spectra, we accounted for context by using a blocked test. This approach addresses the issue where differences in mutation subtype proportions can lead to significant p-values due to variations in the number of specific nucleotide sites, even if the mutation rates are the same across comparison groups. By blocking the test at the trinucleotide context level, we eliminate this source of variation, and the test statistics for all contexts are then combined.

We compared each pair of treatment groups by subsetting the data to include only the mutations from the two groups being compared. For each unique trinucleotide context, Fisher’s exact test was performed to compare mutation counts between the two groups. We then combined the test statistics for all contexts using Fisher’s method for combining p-values. Under the null hypothesis, the sum of these transformed p-values follows a chi-squared distribution with degrees of freedom equal to twice the number of tests. By summing these transformed values, we obtained a combined test statistic. The combined p-value was then computed from this test statistic using the chi-squared distribution. To control for multiple comparisons, we adjusted the combined p-values using the Hochberg method.

### Benchmark dose modelling

Benchmark dose analyses were conducted using the freely-available (https://www.rivm.nl/en/proast) PROAST R-package (version 70.9) in the R programming environment. Dose-response data were analysed using the weighted, four-model average (*i.e.,* exponential, Hill, inverse exponential and log-normal models) approach (EFSA 2022). To prevent overfitting, more complex models with additional parameters were only accepted if the goodness-of-fit exceeded the critical value at P < 0.05. PROAST outputs designate potency (*i.e.,* the BMD) and its 90% confidence interval (*i.e.,* bounded by the BMDL and BMDU values) as the ‘critical effect dose’ (*i.e.,* CED, CEDL and CEDU), respectively. Analyses of *lacZ* mutant frequency (transgenic rodent assay) or mutation frequency response (DS *lacZ* panel) used a benchmark response (BMR) (*i.e.,* the critical effect-size or CES in PROAST notation) of 50% representing interpolation at an effect-size equal to a 50% increase in response relative to the concurrent vehicle control. For model-average bootstrap sampling, 1000 simulations were utilised. All code and PROAST outputs supporting the BMD calculations are available from the BioStudies database (Sarkans et al. 2018) under accession number S-BSST1978.

### Benchmark dose modelling: Application of effect-size theory

To provide more comparable BMD estimates between the transgenic rodent assay (*lacZ*) and DS mouse mutagenesis panel, effect-size theory (Slob 2017) as used to estimate endpoint-specific critical effect-size (ecCES) values that compensated for differences in the two endpoints’ dynamic range (*i.e.,* maximum response) (analyses shown, Supplementary Figure 3). Using the PROAST R-package (version 70.9), the profile likelihood method was used to estimate parameter *c* (*i.e.,* the maximum response of each endpoint) and its 90% confidence interval using the exponential m5 model. As the dose-response data for both the TGR (*lacZ*) and DS endpoints showed sigmoidal relationships with doses high enough to capture the ‘plateauing’ in the response that occurs towards the maximum response, the confidence intervals for the parameter *c* estimates were quite precise in each instance (< 2-fold) despite estimation from single dose-response relationships. For this reason, preliminary ecCES values were estimated from the maximum response estimates in terms ecCES = (*c*^1/8^ -1)*100 (Slob 2017). Whereas the presented analyses demonstrate the concept, analyses of multiple datasets will be required to establish final, *generalisable*, ecCES values that can be recommended for risk assessment purposes. In this work, the goal was to simply to demonstrate the need for the approach, and to obtain BMD estimates for the paired, TGR (*lacZ*) and DS (mutagenesis panel) data that were better-suited for comparison. All code and PROAST outputs supporting the esCES work are available from BioStudies database (Sarkans et al. 2018) under accession number S-BSST1978.

## RESULTS

### NDMA-induced mutagenesis determined by DS is dose dependent in Muta™Mouse

We performed DS on the liver samples from the OECD-compliant TGR mutation assay (Table 1, (Lynch et al. 2024)). In contrast to the TGR study, which provides an indirect measure of mutation at a single artificial transgenic locus, DS was deployed to directly detect mutations by generating DS libraries for two panels; the mouse mutagenesis panel (Supplementary Figure 1) designed to be representative of the wider mouse genome (LeBlanc et al. 2022) and a second *lacZ* panel to sequence mutations that occur only within the bacterial *lacZ* transgene (included to provide a more direct comparison with the TGR *lacZ* mutant data). For the mouse mutagenesis panel an average of 515,025,886 raw reads were generated per sample leading to an average of 1,282 million (1,136 – 1,407 million) informative duplex bases per sample. For the *lacZ* panel an average of 312,888,494 raw reads were generated per sample leading to 1,140 million (794 – 1,347 million) informative duplex bases (Supplementary Table 1). By design, similar numbers of informative duplex bases were obtained from both panels by adjusting the starting amount of DNA in the library preparation to account for the fact that the *lacZ* panel (3,089 bp) is much smaller than the Mutagenesis Panel (48,000 bp) and the *lacZ* gene is present as a multi-copy transgene. The result of this is that, while the number of duplex bases is similar, the two panels have different on-target duplex depths, the average on-target duplex depth for the Mutagenesis Panel is 26,711 (23,663 – 29,320) and for the *lacZ* panel is 275,554 (190,513 – 324,452).

**Table 1.** Study design in male Muta™Mouse treated with NDMA. *TGR study is from Lynch et al., 2024.

| Treatment Group | Dose (mg/kg/day) | Total dose (mg/kg) | Dose regimen | N | TGR* | Duplex Sequencing |  |
| --- | --- | --- | --- | --- | --- | --- | --- |
|  |  |  |  |  |  | LacZ | Mouse Mutagenesis |
| TS Control | 0 | 0 | - | 1 |  | ✓ | ✓ |
| Vehicle Control | 0 | 0 | - | 6 | ✓ | ✓ | ✓ |
| NDMA | 0.02 | 0.56 | Daily | 5 | ✓ | ✓ | ✓ |
| NDMA | 0.07 | 1.96 | Daily | 6 | ✓ | ✓ | ✓ |
| NDMA | - | 5 | Single | 5 | ✓ | ✓ | ✓ |
| NDMA | 0.19 | 5.32 | Daily | 6 | ✓ | ✓ | ✓ |
| NDMA | - | 10 | Single | 4 | ✓ | ✓ | ✓ |
| NDMA | 0.36 | 10.08 | Daily | 6 | ✓ | ✓ | ✓ |
| NDMA | 1.1 | 30.8 | Daily | 5 | ✓ | ✓ | ✓ |
| NDMA | 2.0 | 56 | Daily | 5 | ✓ | ✓ | ✓ |
| NDMA | 4.0 | 112 | Daily | 5 | ✓ | ✓ | ✓ |
| Positive control (ENU) | - | 120 | 3 x Single | 3 | ✓ | ✓ | ✓ |

N number of samples

For the repeat dose regimen, a dose-dependent increase in mutation frequency was observed for NDMA-induced mutation in the *lacZ* panel (Figure 1a and supplementary table 2) with statistically significant increases observed at a total dose of 0.19 mg/kg/day (5.32 mg/kg) and above compared with the vehicle control; there were no significant increases in mutation frequency observed at the two lowest doses of NDMA. A similar dose response was observed across the wider mouse genome using the mutagenesis panel (Figure 1b and supplementary table 2), although background mutation frequencies using the *lacZ* panel were higher than the mutagenesis panel, and the latter exhibited a greater dynamic range in terms of overall response. For both the bacterial *lacZ* and mouse mutagenesis panels, the Lowest-Observed Genotoxic Effect Level (LOGEL) was 0.19 mg/kg/day (total dose 5.32 mg/kg) with a No-Observed Genotoxic Effect Level (NOGEL) of 0.07 mg/kg/day ((total dose 1.96 mg/kg).

There were increases in mutation frequency observed for the two single bolus treatment groups (5 and 10 mg/kg NDMA) using both sequencing panels (Figures 1c and d), yet unlike the TGR *lacZ* mutant frequency (Lynch et al. 2024), these increases were not significantly different statistically from those observed in the repeat dose treatment groups with equivalent total doses. However, some differences in mutation type were seen. NDMA induces mainly C>T transitions (Delker et al. 2008; Shane et al. 2000; Souliotis et al. 1998; Wang et al. 1998). The single bolus dose groups displayed higher T>G mutation frequencies relative to the repeat dose groups in the *lacZ* and Mouse Mutagenesis panels. There were also lower C>T mutation frequencies in the single dose group at 10 mg/kg relative to the repeat dose group, but these were only statistically different in the *lacZ* panel (Supplementary Figure 2). The reason for these differences remains unclear although the DS results are consistent with Haber’s law, which states the incidence and/or severity of a toxic effect (*sic* mutagenesis) depends on total exposure, i.e. exposure concentration (c) rate times the duration time (t) of exposure (i.e., c x t).

### Comparison of DS mutation frequency to TGR *lacZ* mutant frequency dose response data induced by NDMA

The DS mutation frequency data were compared with the *lacZ* gene mutant frequency from the TGR mutation study. The TGR study measures the proportion of *lacZ* genes that harbour a functional mutation whereas Duplex Sequencing directly measures the frequency of all mutations. The *lacZ* mutant frequency results from the TGR mutation assay were highly correlated (P > 0.95) with the DS *lacZ* mutation frequencies (Figure 2a) and Mouse Mutagenesis mutation frequencies (Figure 2b) from across the entire dose range. Notably, DS was able to identify a significant increase in mutation incidence at doses as low as 0.19 mg/kg/day (5.32 mg/kg), a sensitivity not observed with the TGR mutation assay. The enhanced sensitivity of DS, particularly at lower doses, can be partially attributed to the reduced variability in mutation frequency values observed compared to the phenotypic *lacZ* mutant frequencies in the TGR mutation assay: The standard deviation within groups was significantly lower for both DS-panels (Mouse Mutagenesis and *lacZ*) compared to TGR mutation assay (Figure 2c). Moreover, the DS data corrects for clonality (using the min method), a correction that is not possible for the TGR mutation assay without time-consuming Sanger sequencing of mutant clones. When clonality correction was removed from the DS data (max method), the in-group variability was much higher (Figure 2d), and the *lacZ* panel data was comparable to the TGR *lacZ* assay data (P > 0.05, Student’s T-test) but this was not the case with the Mouse Mutagenesis panel (P < 0.05).

### Benchmark dose modelling of DS mutation frequency and TGR mutant frequency data suggest concordant point-of-departure estimations

To determine the mutagenic potency of NDMA in the repeat-dose study, benchmark dose analysis was performed, harnessing data from the complete dose-response relationships for both DS and TGR endpoints. As the dynamic ranges (*i.e.,* maximum response parameter *c* estimates, shown Supplementary Figure 3) were near-identical for the TGR *lacZ* and DS *lacZ* endpoints, BMD_50_ values were estimated and directly compared (Figure 3a-c). Whereas the BMD confidence interval was smaller (*i.e.,* more precise) from the DS (*lacZ*) data (due to tighter within-group variation) the comparison established BMD_50_ values with overlapping confidence intervals (Figure 3c). This shows that the BMD for DS (*lacZ*) was not significantly different to the BMD derived from the TGR (*lacZ*) assay. In other words, using data established from paired tissue samples, the benchmark dose modelling provides supporting evidence for concordance between the DS (*lacZ*) analysis and the traditional transgenic rodent assay. Further preliminary BMD comparisons for DS (mouse mutagenesis panel) and the TGR assay using effect-size theory (Slob 2017) to normalise for differences in dynamic range are presented in Supplementary Figure 3.

**Fig. 3.**
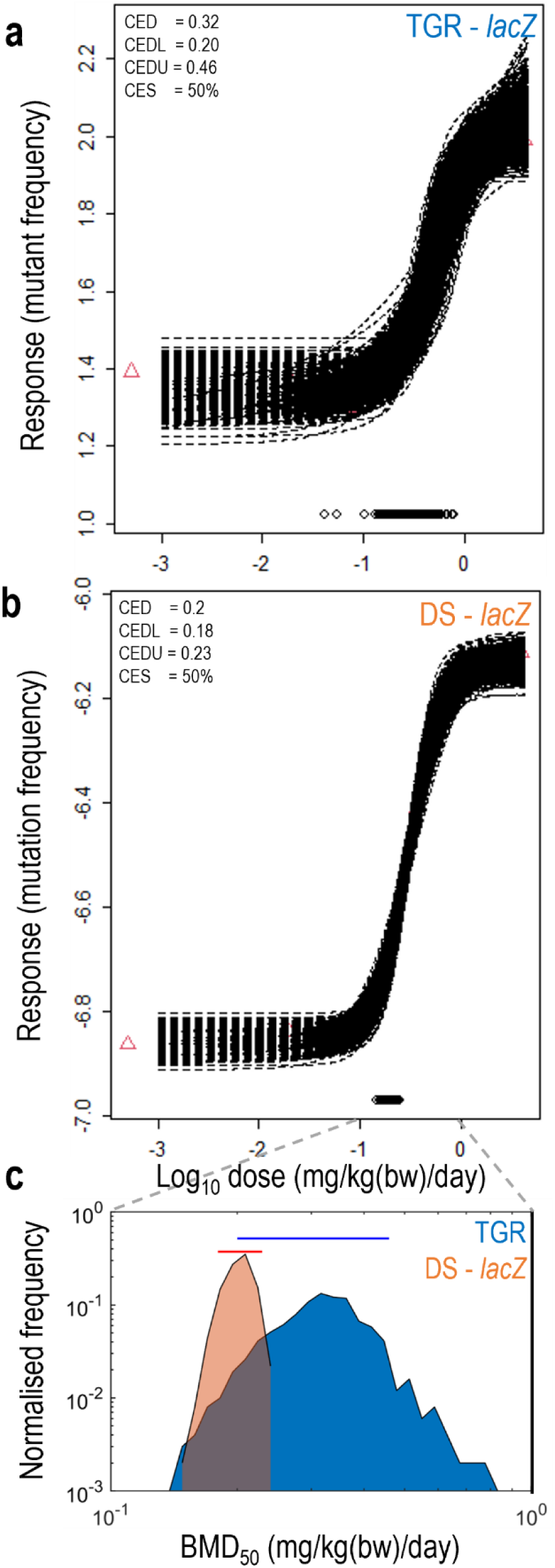
Comparison of benchmark dose estimates from TGR or DS methods. **a/b**, Four-model average, benchmark dose modelling fits to the dose-response data from **a** TGR (*lacZ* transgene) or **b** DS (*lacZ* panel). A fixed critical-effect size of 50% was used for both TGR and DS endpoints and 1000 bootstrap simulations were run. The points clustered along the X-axis show the estimated critical-effect dose (CED) from each bootstrap run. The CEDL and CEDU values represent the lower and upper 90% confidence interval of the CED, (respectively). **c**, Distributions (histograms) and 90% confidence intervals (horizontal lines) of the critical-effect dose estimates. Whereas the CED estimates from the TGR and ecNGS methods overlapped, ecNGS yielded a narrower distribution (i.e., greater precision) positioned at the left-end (i.e., more sensitive) of the results from the TGR method.

### Single base substitution spectra induced by NDMA in Muta™Mouse

Mutant *lacZ* plaque forming units were not sequenced in the original TGR mutation assay and therefore concurrent sequence context mutation type data from *lacZ* mutant clones were not available. DS data contains comprehensive information about the location and type of each mutation detected, and this was used to generate single base substitution spectra. For both the *lacZ* panel (Figure 4a) and Mouse Mutagenesis panel (Figure 4b), the mutations observed upon high dose NDMA treatment were predominantly C>T transitions, as expected based on the known mechanism of action for NDMA, and consistent with other published TGR studies (Armijo et al. 2023; Delker et al. 2008; Shane et al. 2000; Souliotis et al. 1998; Wang et al. 1998). A different spectrum was seen with the positive control ENU, with an enrichment for T>A and T>C changes consistent with previous data (Smith-Roe et al. 2023). Where the two DS panels differ is in the spectra for vehicle control samples. In the mutagenesis panel a range of mutation types was observed but, for the *lacZ* panel, the C>T transition mutation was predominant.

**Fig. 4.**
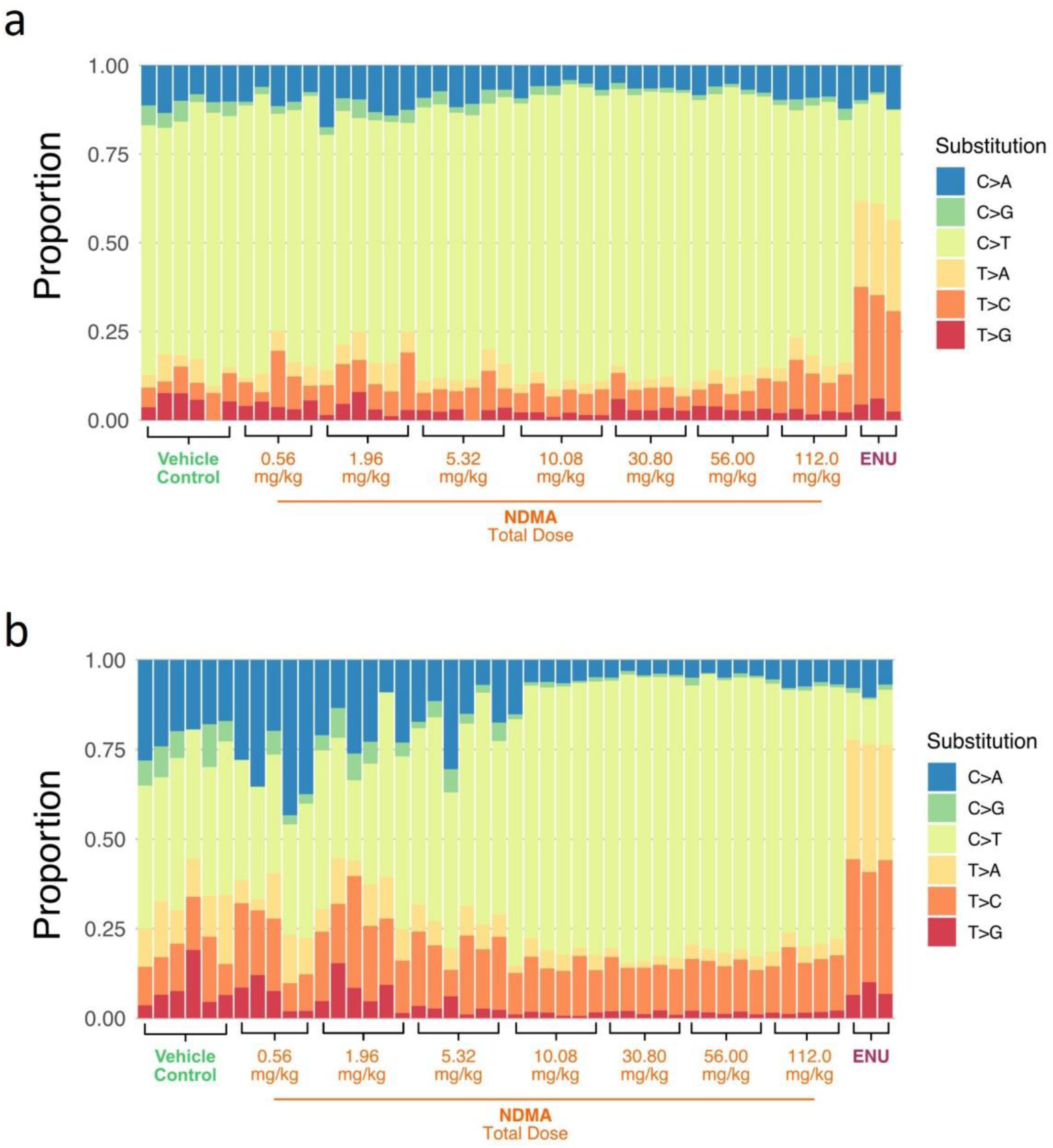
Proportions of single base substitution types in Muta™Mouse liver following NDMA exposure using Duplex Sequencing. Stacked bar plots display the proportions of single base substitutions across varying doses of NDMA and a positive control (ENU) using Duplex Sequencing of **a** the *lacZ* panel and **b** mouse mutagenesis panel. The colour key to the right of the panels identifies the substitution types.

The mutagenesis panel, by design, contains a trinucleotide abundance profile very similar to the wider mouse genome (Figure 5a). Unsurprisingly, the prokaryotic *lacZ* gene has a different trinucleotide abundance profile to either the mutagenesis panel or mouse genome (Figure 5b). Therefore, it was important to consider whether the differences in trinucleotide context abundance impacted the mutation frequency and single base substitution spectra obtained by DS. The data from the mutagenesis panel was used to generate a predicted mutation frequency and a predicted single base substitution spectrum for the *lacZ* panel: the number of mutations and the number of duplex bases observed at each trinucleotide context in the data from the mutagenesis panel was multiplied by the relative trinucleotide context abundance between the *lacZ* and mutagenesis panel. The predicted mutation frequency for *lacZ* became more closely aligned with the observed mutation frequency for this locus (Figure 5c). This increased similarity was consistent across all dose levels, with the correlation between the panels significantly improving after adjusting for trinucleotide context differences (Figure 5d). Moreover, the adjustment also explains the higher frequency of C>T substitutions observed in the *lacZ* panel’s vehicle control data, with the predicted values very similar to those observed in the actual data from the *lacZ* panel (Figure 5e). This implies that the observed differences between these panels are largely the result of differences in genomic composition between the bacterial *lacZ* gene and the DNA sequences that make up the mouse mutagenesis panel, underlining the importance of genomic composition on mutation frequency and spectra.

**Fig. 5.**
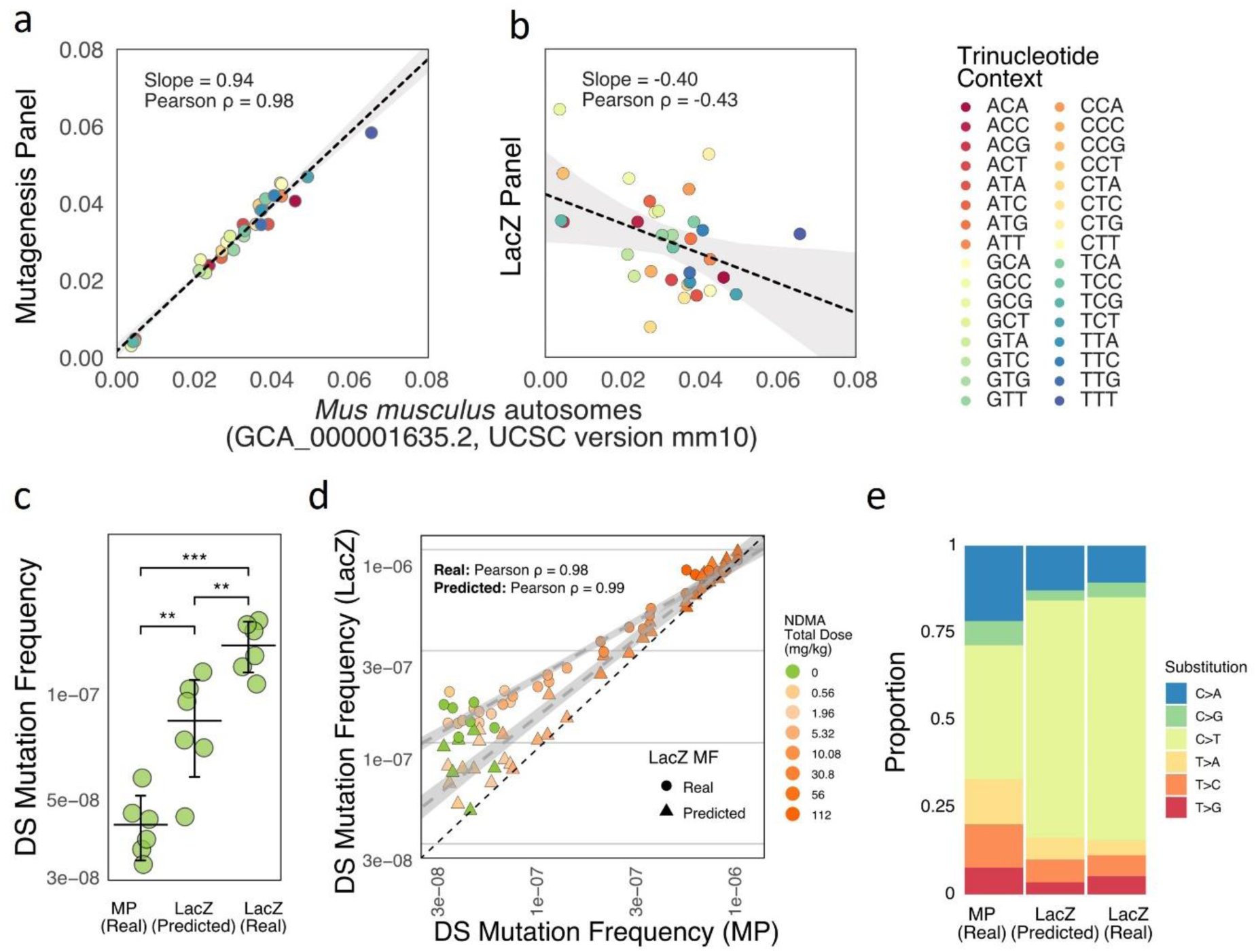
Correction for trinucleotide context content to predict mutation frequency and substitution types in Muta™Mouse liver. Correlation of trinucleotide context abundance from **a** the Mutagenesis panel and **b** the *lacZ* panel with endogenous mouse genome. The colour key identifies the trinucleotide context for each data point. **c** Comparison between DS Mutation Frequency using mutagenesis panel (MP) and for real data from the *lacZ* panel (*lacZ* Real), as well as MP data corrected for trinucleotide context abundance differences between the panels (*lacZ* Predicted). **d** Correlation of DS mutation frequencies between real and predicted data for the *lacZ* panel, with NDMA total doses indicated in the colour key. **e** Proportions of single base substitution types from the Mutagenesis Panel and for real and predicted data *lacZ* panels. The colour key to the right of the panels identifies the substitution types. Statistical significance is denoted by *** (p < 0.001) and ** (p < 0.01).

### Trinucleotide spectra induced by NDMA reveals a characteristic signature

DS data contains both mutation type and mutation location information, allowing the generation of 96-channel trinucleotide spectra. In both the *lacZ* (Figure 6a) and mouse mutagenesis (Figure 6b) panels, consistent mutation patterns were observed, dependent on NDMA dose. The spectra for 28-day NDMA total doses at or below 0.07 mg/kg/day, as well as vehicle controls, primarily comprised C>T substitutions at CpG sites. This trinucleotide spectrum is reminiscent of the SBS1 clock-like signature (Cosine Similarity between SBS1 and Vehicle Control = 0.91) recorded in the COSMIC Database (Tate et al. 2019). The proposed aetiology is spontaneous cytosine deamination and has previously been observed by DS in the TGR assay (LeBlanc et al. 2022). At higher total NDMA doses, a distinctive trinucleotide mutation pattern emerged. Starting from approximately 0.19 mg/kg/day, this pattern was increasingly characterized by C>T substitutions in 5’-N–C–Pyrimidine –3’ contexts (Figure 6) and was comparable to the SBS11 signature in COSMIC (Cosine Similarity between SBS11 and 4 mg/kg/day NDMA = 0.89). The SBS11 signature is associated with alkylating agents and therefore aligns with the known mechanism of NDMA induced mutagenesis. Trinucleotide spectra correct for trinucleotide context abundance and so direct comparisons can be made between the *lacZ* and mouse mutagenesis panels. The mutation patterns showed significant alignment, underscoring a consistent mutational response to the treatments across the regions sampled.

**Fig. 6.**
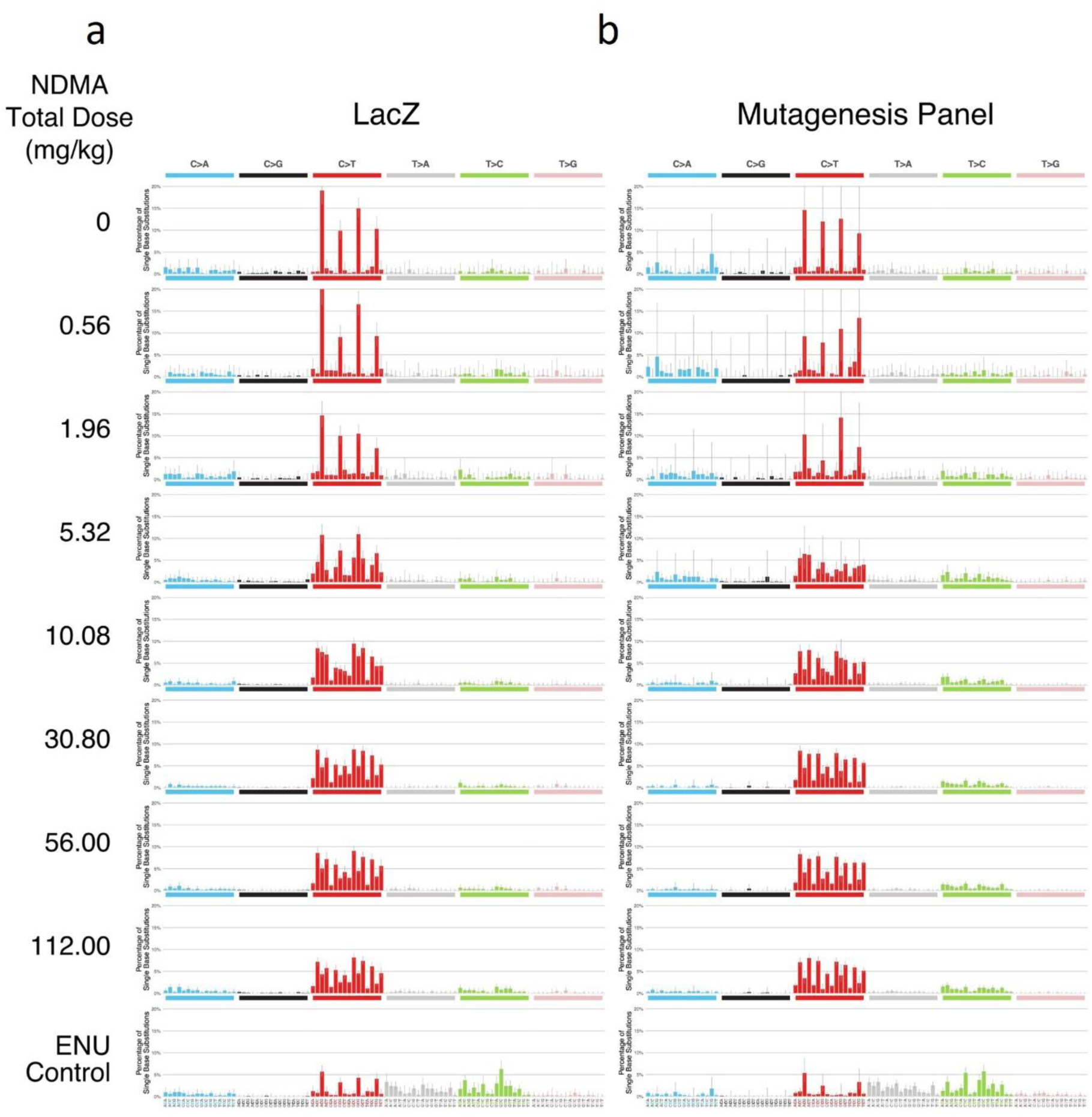
Trinucleotide mutation spectra induced by NDMA of Muta™Mouse liver using Duplex Sequencing of *lacZ* and Mutagenesis panel. Mutation spectra using **a** the *lacZ* panel and **b** the mutagenesis panel at various total doses of NDMA compared to the ENU control. Each panel represents the pooled mutation spectrum across animals at a specific dose, with mutations categorized by trinucleotide context. The x-axis in each panel represents the sequence context of the mutated base, while the y-axis represents the frequency of the mutations, corrected for sequencing depth at the relevant trinucleotide context abundance.

The trinucleotide mutation spectra for the mouse mutagenesis panel were compared mathematically using cosine similarity (Figure 7a). Unsupervised hierarchical clustering of cosine similarity scores revealed 3 distinct clusters and remarkable concordance between treatment groups with DS at the trinucleotide spectrum level. The vehicle control spectrum was similar to the two lowest doses of NDMA and SBS1; these data directly support the existence of a NOGEL for NDMA in the Muta™Mouse study. At doses of 0.19 mg/kg/day (LOGEL) and above, the cosine of similarity values were similar to one another and the SBS11 signature, and moreover, the mutational signature reported by Armijo et al. in another mouse strain and observed at doses of NDMA commensurate with the higher doses used in the current study (Armijo et al. 2023). In contrast, the mutation spectrum induced by ENU was similar to that of ENU from the Signal Database (Srinivasan et al. 2021) and unlike the vehicle control or NDMA. Statistical comparison of NDMA spectrum data between the Muta™mouse treatment groups demonstrated that doses of 0.19 mg/kg/day and above were statistically distinct from the spectra generated by the vehicle controls and two lowest doses (Figure 7b). These observations provide robust orthogonal evidence for a detectable genotoxic effect occurring only at doses of 0.19 mg/kg/day NDMA and above and substantiate the veracity of the mutation frequency findings.

**Fig. 7.**
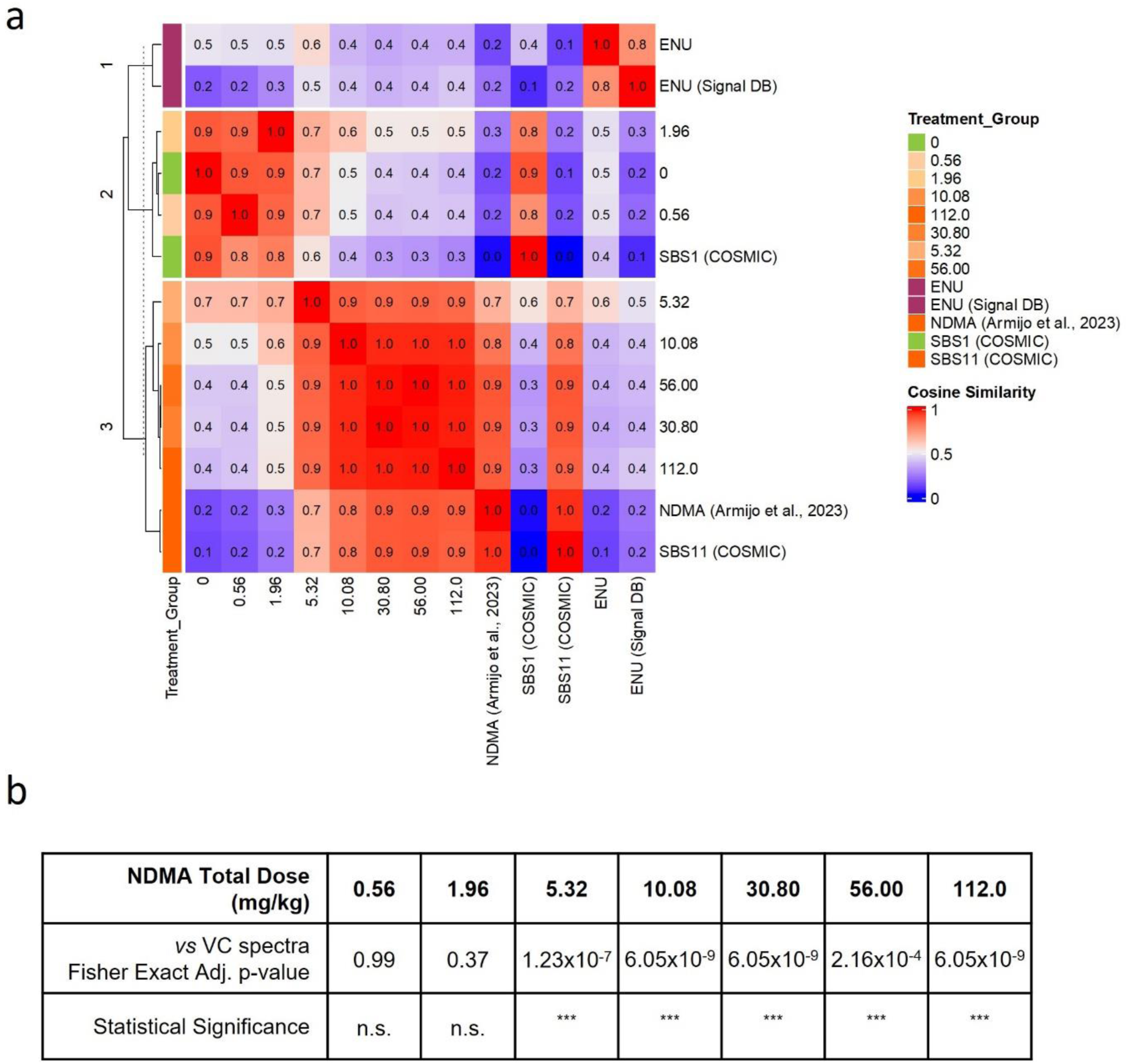
Comparison between trinucleotide spectra across treatment groups using Duplex Sequencing of the mutagenesis panel. **a** Heatmap illustrating the cosine similarity between trinucleotide mutation spectra of different NDMA total dose groups, controls and several reference mutational signatures. Comparative reference mutational signature includes NDMA from Armijo et al., 2023, ENU from the Signal Database and SBS1 and SBS11 from COSMIC (v3.4). The colour scale represents cosine similarity values, ranging from 0 (low similarity) to 1 (high similarity). Hierarchical clustering on the row groups the treatment conditions based on similarity. **b** Statistical analysis table showing comparison between trinucleotide spectra of NDMA-treated groups and the vehicle control (VC) spectra. The table includes Fisher Exact adjusted p-values and statistical significance annotations (n.s. for not significant, *** for p < 0.001) for each dose group.

## DISCUSSION

This study utilised DS, a highly sensitive and error-corrected DNA sequencing method, using a TGR transgene panel (*lacZ*) in addition to the mouse mutagenesis panel, allowing separation of results from genome-representative DS from the classic TGR assay to infer what differences in those are due to the assays versus the different genomic context being examined. DS detected NDMA-induced mutagenesis *in vivo* and demonstrated its capability to identify genotoxic effects at doses undetectable by standard TGR mutation assays and with significantly higher precision than TGR at all dose levels. The study found a lowest observed genotoxic effect level (LOGEL) for NDMA at 0.19 mg/kg/day in a 28-day repeat dose study design and defined a no observed genotoxic effect level (NOGEL) at 0.07 mg/kg/day. Recently, the presence of liver NOGELs for mutagenesis in rodent 28-day TGR mutation assays been observed with other N-nitrosamines, namely, N-nitrosodiethylamine (NDEA), N-nitrosopiperidine (NPIP) and a novel N-nitrosamine impurity identified in sitagliptin-containing products, 7-nitroso-3-(trifluoromethyl)-5,6,7,8-tetrahydro-[1,2,4]triazolo-[4,3-a]pyrazine (NTTP) (Bercu et al. 2023; Powley et al. 2024; Zhang et al. 2024). However, only the NDEA studies were confirmed by DS (Bercu et al. 2023; Zhang et al. 2024). In this study, at higher doses, the DS mouse mutagenesis panel reported an approximately 13.6-fold increase in mutation burden following NDMA treatment whereas only an approximately 4.8-fold increase was observed in the corresponding TGR mutation assay. Similarly, differences in DS fold-change compared with TGR for nitrosamines have been reported elsewhere (Bercu et al. 2023; Zhang et al. 2024). Interestingly, comparison of DS analysis at the *lacZ* locus (approximately 5.2-fold) was more closely aligned with the TGR *lacZ* mutant frequency (approximately 4.8-fold), perhaps suggesting that the varying fold-change differences compared with the TGR assay are more driven by genomic locus and nucleotide composition than the technique used.

DS also provided detailed mutation type and location data, which revealed that C>T transitions were the predominant induced mutation type for NDMA at higher doses, aligning with its known mechanism of action involving the formation of O6-methylguanine adducts and previous reports of NMDA-induced mutations in rodents (Delker et al. 2008; Loechler et al. 1984; Shane et al. 2000; Souliotis et al. 1998; Wang et al. 1998). The mutation spectra from the higher doses of NDMA were similar to the COSMIC SBS11 signature, which is associated with mutations resulting from DNA alkylation, and differed from the clock-like mutation signatures observed in vehicle controls and in the two lowest dose NDMA treatments. Interestingly, whilst the results from this study were similar to those observed in another *in vivo* study in mouse (Armijo et al. 2023), they were different from the mutation type observed in a human liver model where T>C was the predominant mutation type observed (Seo et al. 2024). These observations highlight the potential influence of exposure, metabolism, species and repair mechanisms in different biological test systems. The results in Muta™mouse are consistent with the notion that at low doses of NDMA, which are more commensurate with those anticipated from exposures to trace impurities in the environment and some medicines, there are no detectable increases in mutation frequency above that seen over background. Interestingly, the relative risk for tumorigenesis has been reported for NDMA following extensive lifetime studies in the rat (Peto et al. 1991a; Peto et al. 1991b) and showed no observable increases in liver tumour burden compared with controls at lower doses. Moreover, the adverse outcome pathway for DNA alkylation leading to cancer (AOP 149) recognises DNA mutation as a necessary and essential key event leading to increased cancer risk and, therefore, if mutation is absent then there can be no concern for tumorigenesis. These observations have important implications for the risk assessment of NDMA impurities in medicines and for the carcinogenicity risk assessment.

DS exhibited higher precision and lower variability in mutation frequency values across individual animal samples compared to the same comparable animal TGR *lacZ* mutant frequencies. This in part was due to the ability of DS to correct for clonal amplification. Similar observations have been reported elsewhere with procabazine (Dodge et al. 2023). In addition, by being able to identify the exact position and mutation type, DS is able to recognize germline variants in the transgene which may or may not be detectable by TGR. DS also observes all mutations, silent and non-silent, not just those that have a consequence on the function of *lacZ*, which may lead to bias in individual animals and therefore may introduce additional variability into TGR assay read out. The ability to correct for clonal amplification, reduce variability, and generate a more accurate estimate of induced mutation frequency *in vivo* represents a significant advantage for using DS over the standard TGR assay. Indeed, DS experiments could be refined using reduced animal numbers per treatment group thereby supporting 3Rs initiatives whilst retaining the statistical power expected by regulatory guidelines for mutagenesis studies *in vivo* (Esina et al. 2024). In addition, the BMD confidence intervals calculated from the DS or TGR (*lacZ*) data were overlapping, supporting concordance between the two methods. BMD analysis has been used to compare DS and TGR dose-response data before (Bercu et al. 2023; Dodge et al. 2023; LeBlanc et al. 2022; Zhang et al. 2024), with three of these four studies reporting discordance between the BMDs defined using TGR or DS data. However, these reports compared TGR data from a transgene to DS data from the mutagenesis panel, whereas in this study, both the TGR assay and DS analysed the *lacZ* transgene. Indeed, we also found that the BMD confidence intervals did not overlap when comparing TGR (*lacZ*) and DS (mutagenesis panel) using BMD_50_ (Supplementary Figure 3). However, analyses of the dose-response data indicated differences in the inducible ranges (i.e., maximum responses) of these two endpoints that are not accounted for by the simple BMD_50_ comparison. Preliminary analyses demonstrating the application of effect-size theory (Slob 2017) to correct for these differences are presented in Supplementary Figure 3. Interestingly this approach restored the overlapping BMD confidence intervals seen for the *lacZ* analyses demonstrating the importance of follow-up work to establish appropriate critical effect sizes for DS endpoints using large datasets containing many dose-response relationships. At the current time, TGR studies are resource intensive and study slots are in limited supply. These results highlight the possibility of running mutagenicity studies in wild-type animals, with integration into repeat dose toxicology studies, thereby further reducing animal use in non-clinical safety assessment and adding to the 3Rs advantages of DS.

This study compared mutation events between the TGR transgene (*lacZ*) and a mouse mutagenesis panel designed to represent the whole mouse genome. While there was a high correlation in NDMA-dose responses between the two, the DS Mutagenesis panel showed a greater dynamic range compared to the TGR *lacZ* assay. Moreover, the NDMA single base substitution profile revealed by the mutagenesis panel included differing proportions of mutation types in NDMA treated samples compared to vehicle controls, including highly significant increases in the proportion of C>T transition mutations. In contrast, the frequencies of C>T transitions in vehicle controls and NDMA treated samples were similar for the *lacZ* panel, resulting in samples not clustering by dose with hierarchical clustering (Supplementary Figure 4). Interestingly, a similarly static global SBS profile was observed in a Big Blue TGR mutation assay (Shane et al. 2000). However, differences were noted in that most of the mutations in the vehicle controls were at CpG sites whereas NDMA SBS was shifted away from CpG locations (Shane et al. 2000). This is consistent with observations from the current study (Figure 6). The reason for these differences between the mutagenesis and *lacZ* panels is likely due, in part, to differences in the sequence of the two panels (Figure 5). Whilst sequence difference alone might explain the differences observed in the single base substitution spectra, it does not fully explain the differences in mutation frequency. The *lacZ* transgene is of bacterial origin and transgenic reporter genes are known to have higher mutation rates than endogenous genes (Lambert et al. 2005). Indeed, such variation in not only restricted to transgenic loci. Analysis of the 20 regions sampled in the DS mutagenesis panel show that genic regions have lower mutation frequencies than intergenic regions (Dodge et al. 2023), and this difference was proposed to be because of transcription coupled repair. The bacterial transgenes in TGR models are not transcribed so it is possible that this higher mutation frequency is due to the absence of transcription coupled repair. It is also possible that other factors such as methylation status, sub-nuclear location or heterochromatin/euchromatin play a contributing role.

The current study identifies several advantages for DS. The ultra-low error-rate allows direct quantification of mutation frequency in genomic DNA, without need of transgenic animals. *In vivo* mutation frequency can be determined using a panel with representative nucleotide composition, rather than a non-representative bacterial transgene, and all mutations are detected whether they are functional or silent, removing selection bias. DS provides information on mutation type and location allowing the removal of jackpot mutations and reducing in-group variability, whilst trinucleotide spectra can be generated to enhance mechanistic insights. The establishment of overlapping BMD confidence intervals also suggests concordance between TGR and DS in terms of PoD estimations. Collectively, these advantages support the use of ecNGS approaches as robust tools for *in vivo* mutagenicity testing. ecNGS has powerful implications for reducing animal use, improving experimental efficiency and the provision of more information to help enable robust regulatory assessments.

## Acknowledgements

We would like to thank Zena Keig-Shevlin, Victoria Brown and other supporting LabCorp staff for conducting the MutaMouse study which included dosing of animals, collection of tissues and scoring the transgene mutation assay.

## Supplementary Information

Submitted in three separate files are four supplementary figures in .pdf format and two supplementary data files in .xlsx format

**Supp. Fig. 1.**
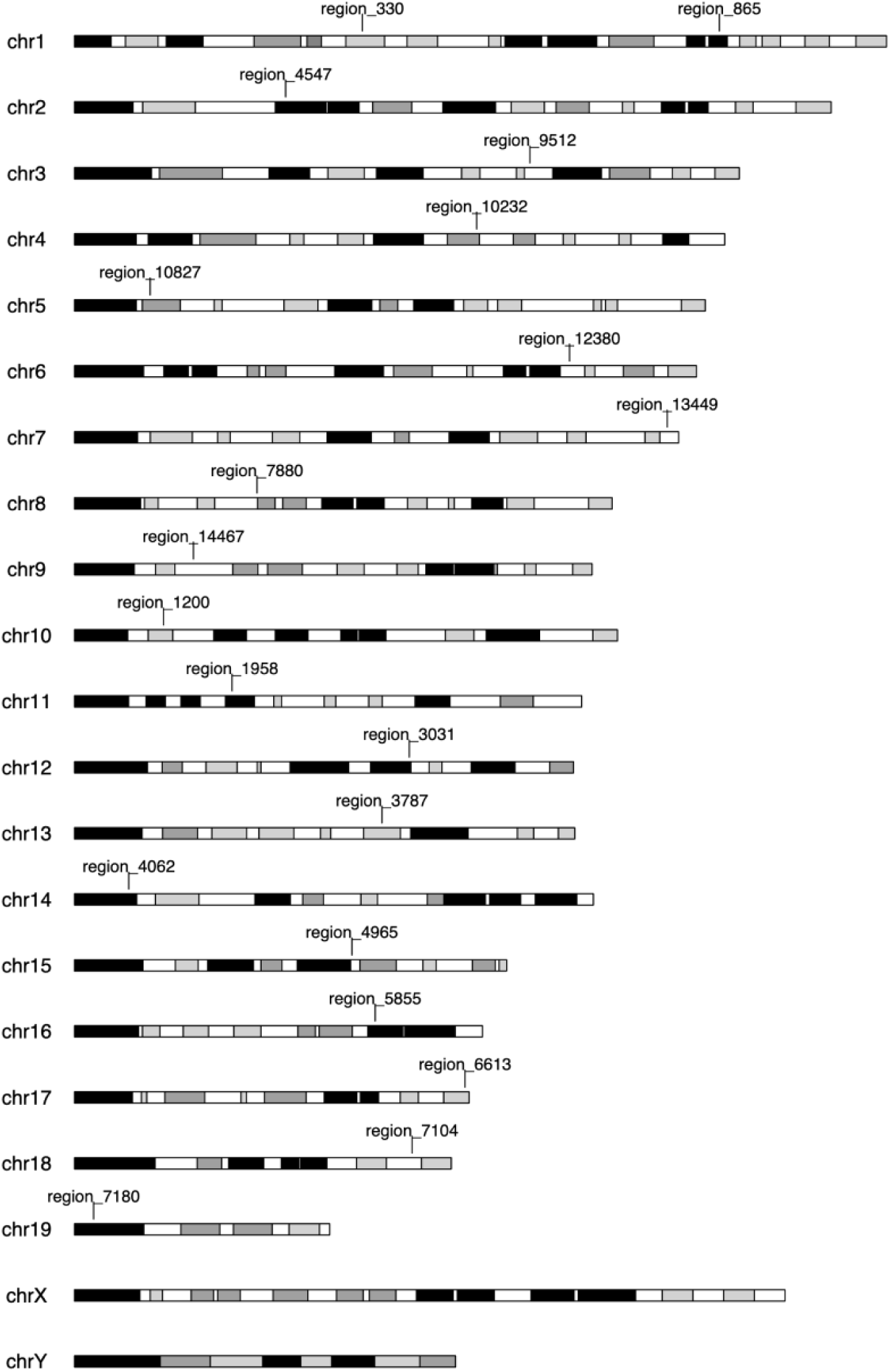
Karyotype diagram of Mutagenesis panel. The mouse mutagenesis panel consists of 20 regions, each 2400 base pairs in length, from across the 20 chromosomes, giving a total panel size of 48 kb. The panel consists of both genic and non-genic targets. Not all genic targets are contained within the coding sequence of the gene and not all genes will be expressed in all tissues

**Supp. Fig. 2.**
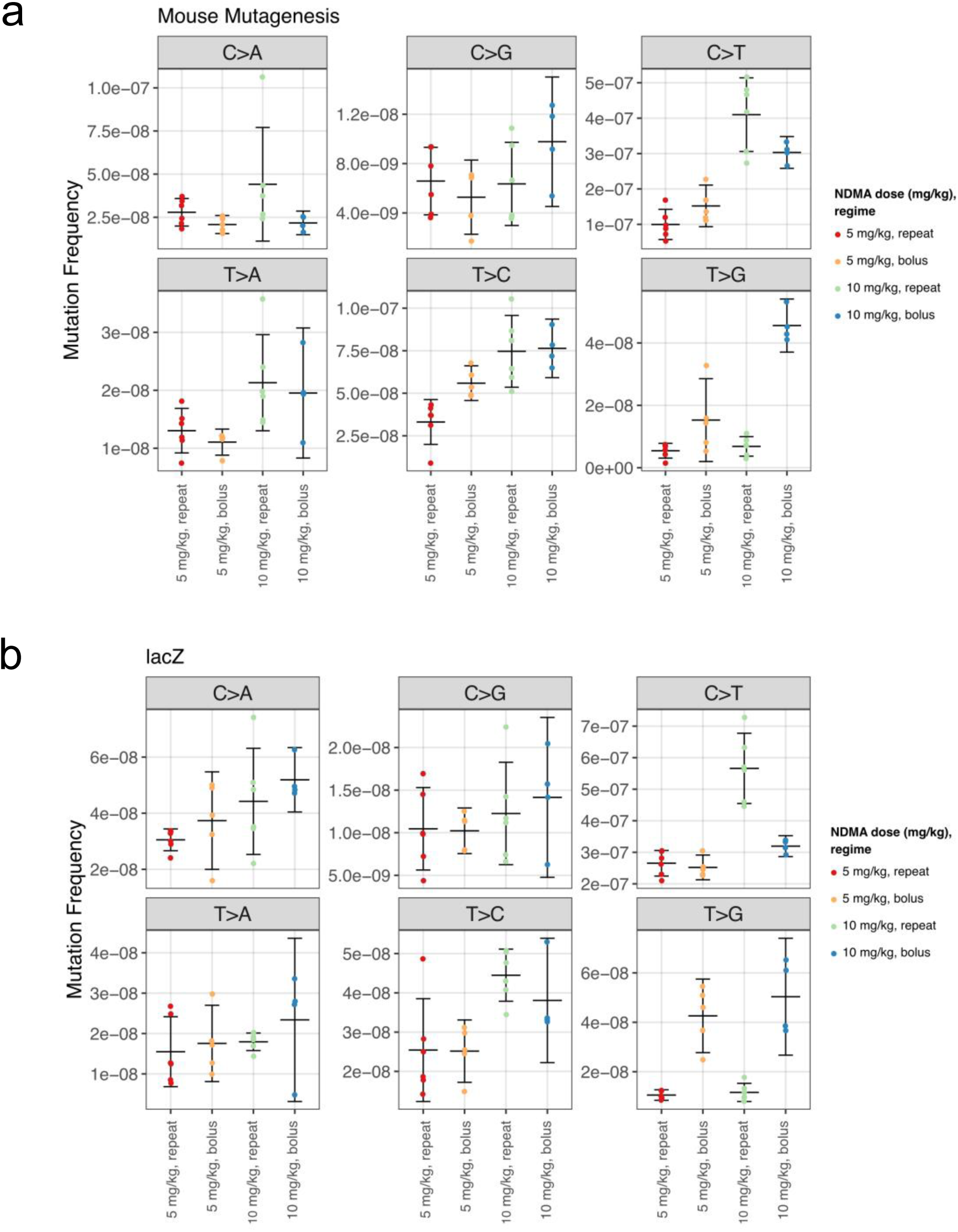
Mutation frequencies of single base substitution types in Muta Mouse liver following NDMA exposure as a function of dose regimen. Each graph represents simple substitution-specific mutation frequency using the **a** *lacZ* panel and **b** Mouse Mutagenesis panel. Error bars represent 95% confidence intervals calculated using a t-distribution

**Supp. Fig. 3.**
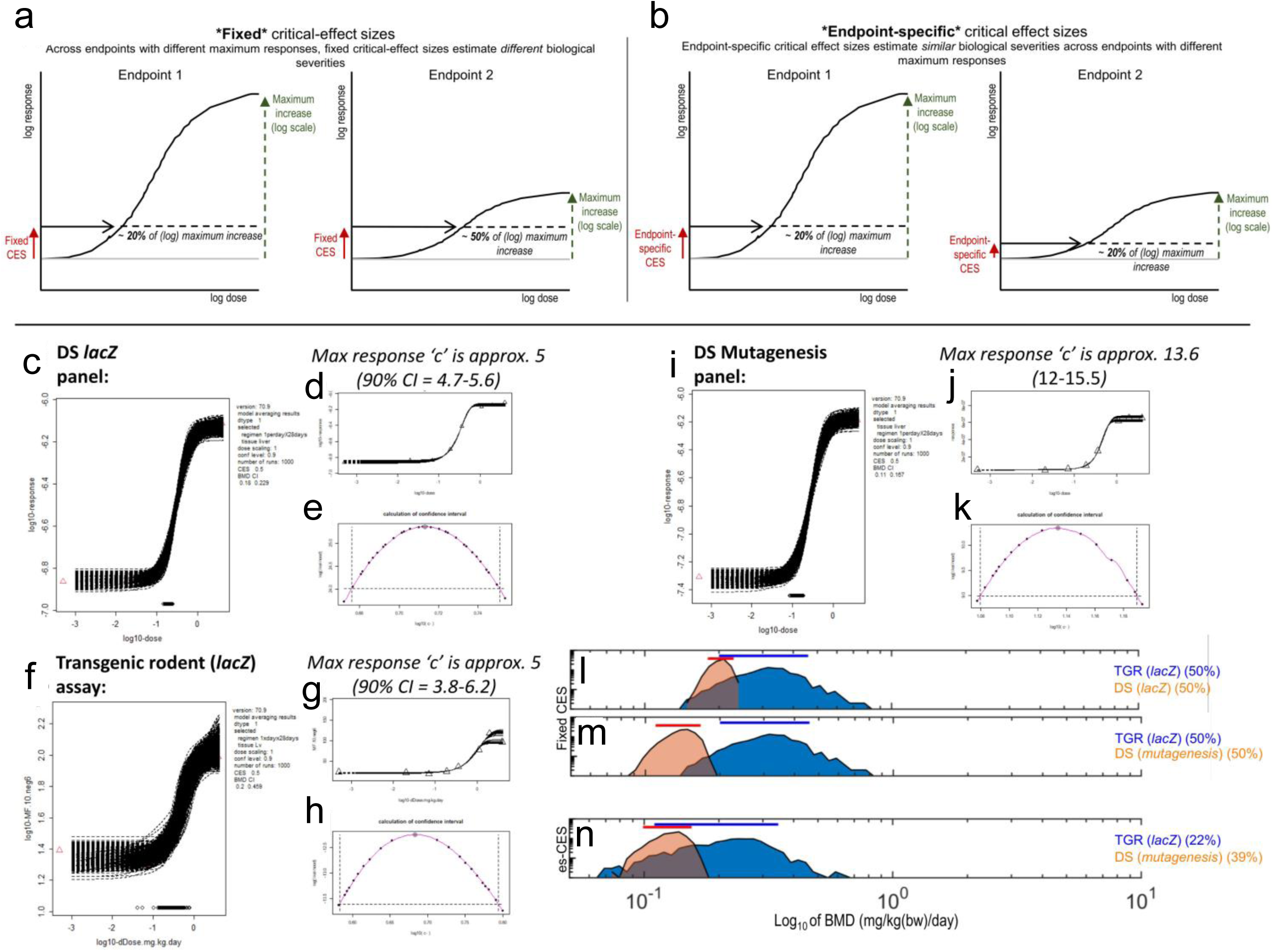
Application of effect-size theory to derive comparable benchmark dose estimates from TGR and ecNGS data. **a/b** schematic explanation of effect-size theory demonstrating the need for endpoint-specific critical-effect sizes (adapted from Beal et al., 2023). **a** When considering two endpoints with different inducible ranges, the use of the same (fixed) critical-effect size (CES) (i.e., relative increase compared to the background response) leads to the determination of critical-effect doses (CEDs) for responses that constitute different proportions of each endpoint’s maximum. In essence, for an endpoint with a large maximum fold increase in inducible response, the magnitude of the response at the CES constitutes only a small fraction of the maximum response inducible for that endpoint (i.e., small effect severity). In contrast, for an endpoint with a smaller maximum fold increase in response, the magnitude of the response at the same CES constitutes a much greater proportion of the maximum for that endpoint (i.e., larger effect severity). For this reason, the CEDs are poorly suited for ‘apples-to-apples’ comparison as the severity of the effects measured are fundamentally different. **b** By scaling the CES to each endpoint’s maximum fold-change by using different, ‘endpoint-specific’ CES values for each endpoint, the magnitude of the response at the critical effect-dose represents equivalent fractions of the maximum fold change for those endpoints (i.e., more comparable effect severities). Importantly, failure to ‘normalise’ effect sizes to the maximum inducible range of the endpoints under comparison will yield critical effect-dose estimates that are unsuitable for direct comparison. **c-n** Application of effect-size theory to TGR and ecNGS dose-response relationships for NDMA. For the TwinStrand *lacZ* panel and the TGR (*lacZ* transgene) endpoints, **d-e/g-h** profile likelihood estimates for model parameter c (maximum response) estimated similar inducible ranges of about ∼5-fold for both endpoints. Therefore, for these two endpoints, CED estimates generated using the same CES are comparable. **l** Shows the 90% CED confidence intervals (lines) and CED distributions (histograms, 1000 bootstrap samples) using fixed CES values of 50%. The CED estimates are observed to overlap, albeit with the ecNGS estimates positioned at the left-end (i.e., more sensitive) of the results from the TGR method. **i-k**, In contrast, a larger inducible range (∼ 14) was estimated from the dose-response data from the TwinStrand Mutagenesis panel. **m** This time, comparing the CED estimates using a fixed CES of 50% established a ∼3-fold lower BMD estimate from the ecNGS data with no overlap in the critical effect-dose confidence intervals (lines). **n** Interestingly, applying effect-size theory and using endpoint-specific critical-effect sizes reconciled these differences, providing overlapping CED distributions in a manner analogous to that observed for **l** TGR and the TwinStrand *lacZ* panel. Combinatorially the analyses presented in this Figure demonstrate the need to consider the dynamic range of the endpoints under comparison when deriving and comparing BMD estimates. Access to a larger cohort of dose-response data from both ecNGS and TGR would establish greater confidence in the apparent similarities in the CED estimates observed between ecNGS and TGR. Whereas only single datasets are available here, the study design is such that the sigmoidal shape of the dose-response is well captured by the employed dose-groups and the maximum response size (parameter c) can still be estimated with some precision (i.e., evidenced by the small 90% confidence intervals for the (**d/g/j**) parameter c estimates). In short, whereas analyses of more datasets are needed, this Figure demonstrates the concept and provides important preliminary data

**Supp. Fig. 4.**
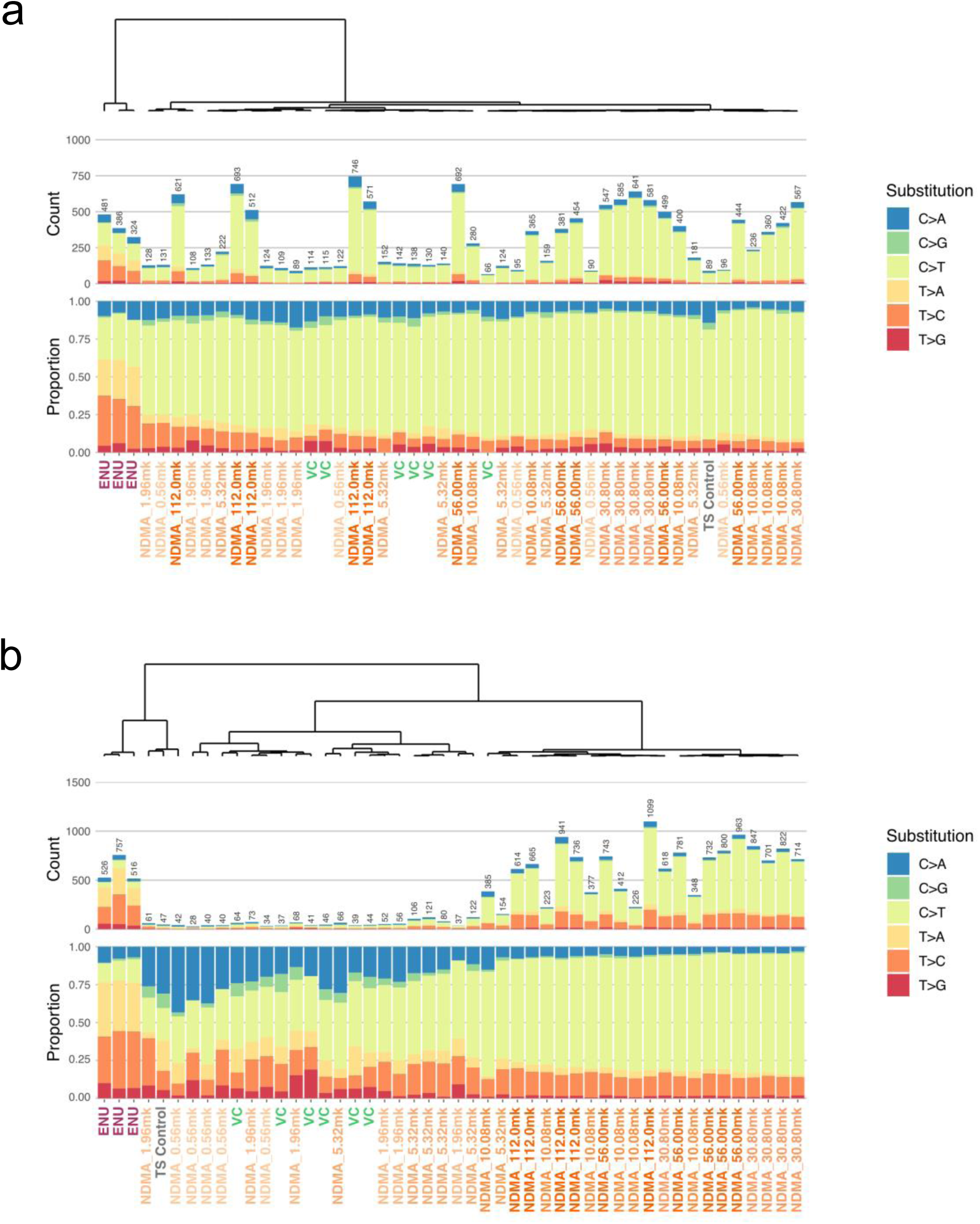
Unsupervised hierarchical clustering of proportion of single base substitution types in Muta Mouse liver following NDMA exposure using Duplex Sequencing. **a+b** Stacked bar plots display the proportions of single base substitutions across varying doses of NDMA and a positive control (ENU) using Duplex Sequencing of the *lacZ* panel (a) and Mouse Mutagenesis panel (b). Dendrograms above each plot represent the results of unsupervised hierarchical clustering using Ward’s D2 method and the cosine similarity metric. The color key to the right of the panels identifies the substitution types

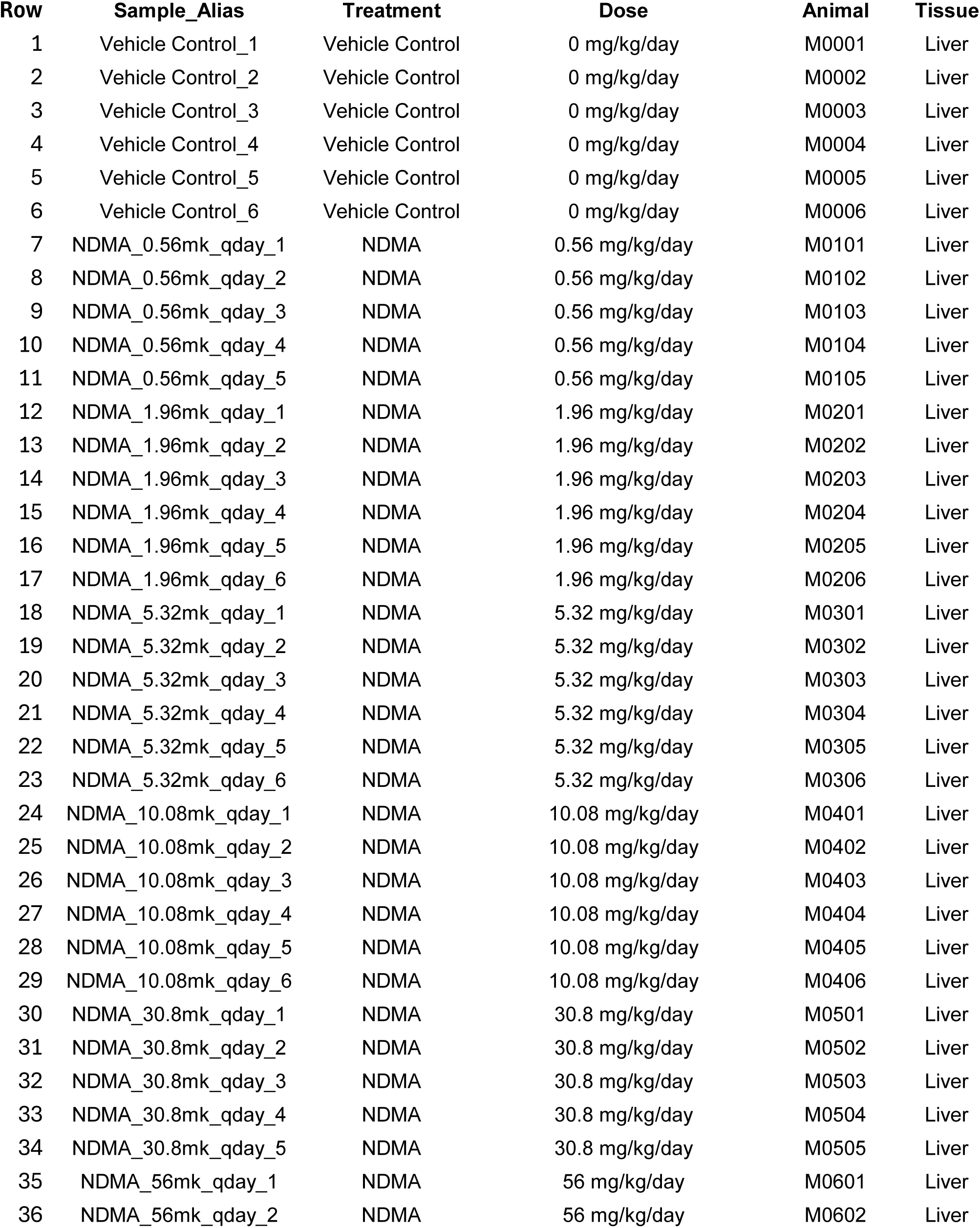
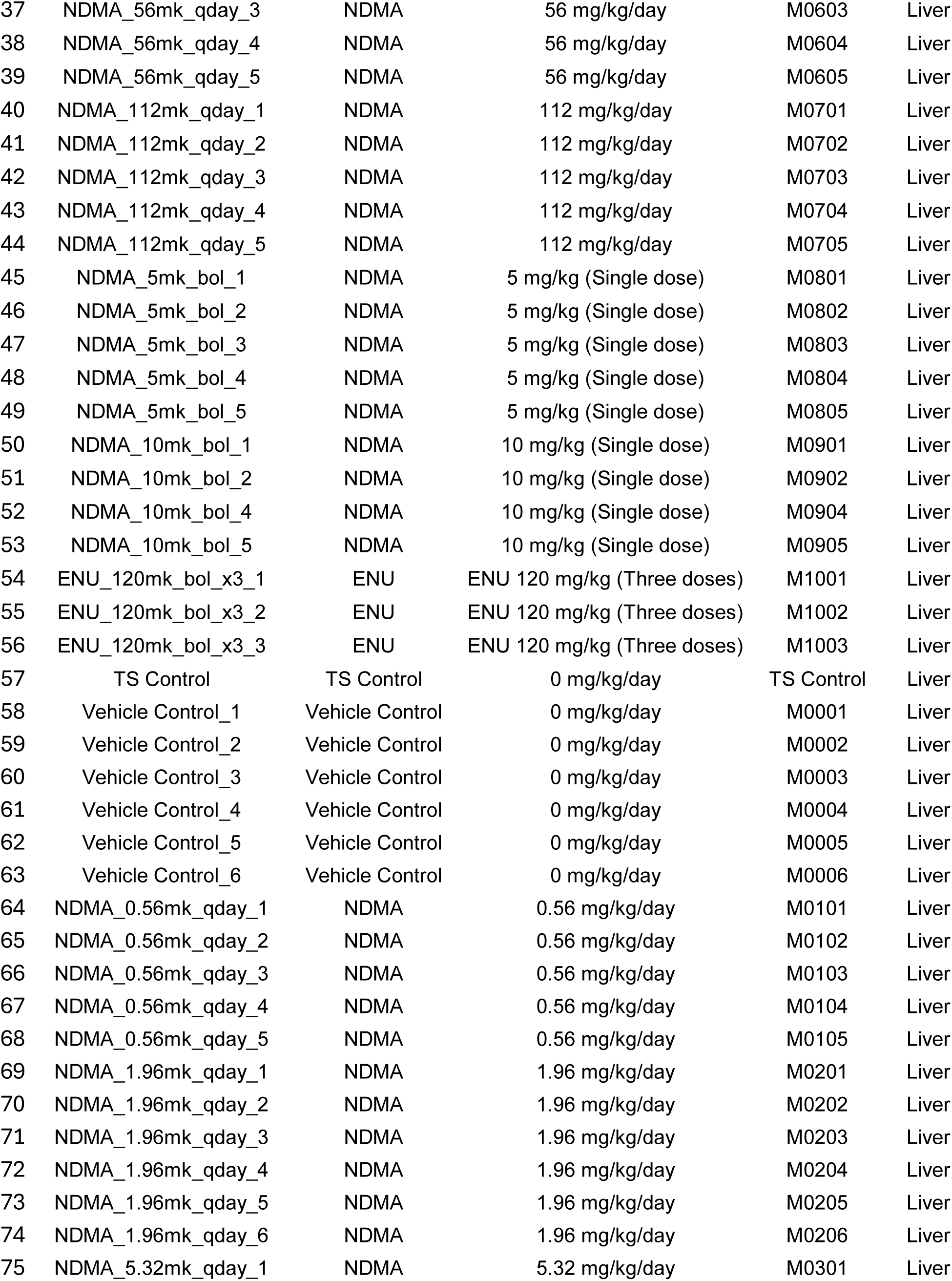
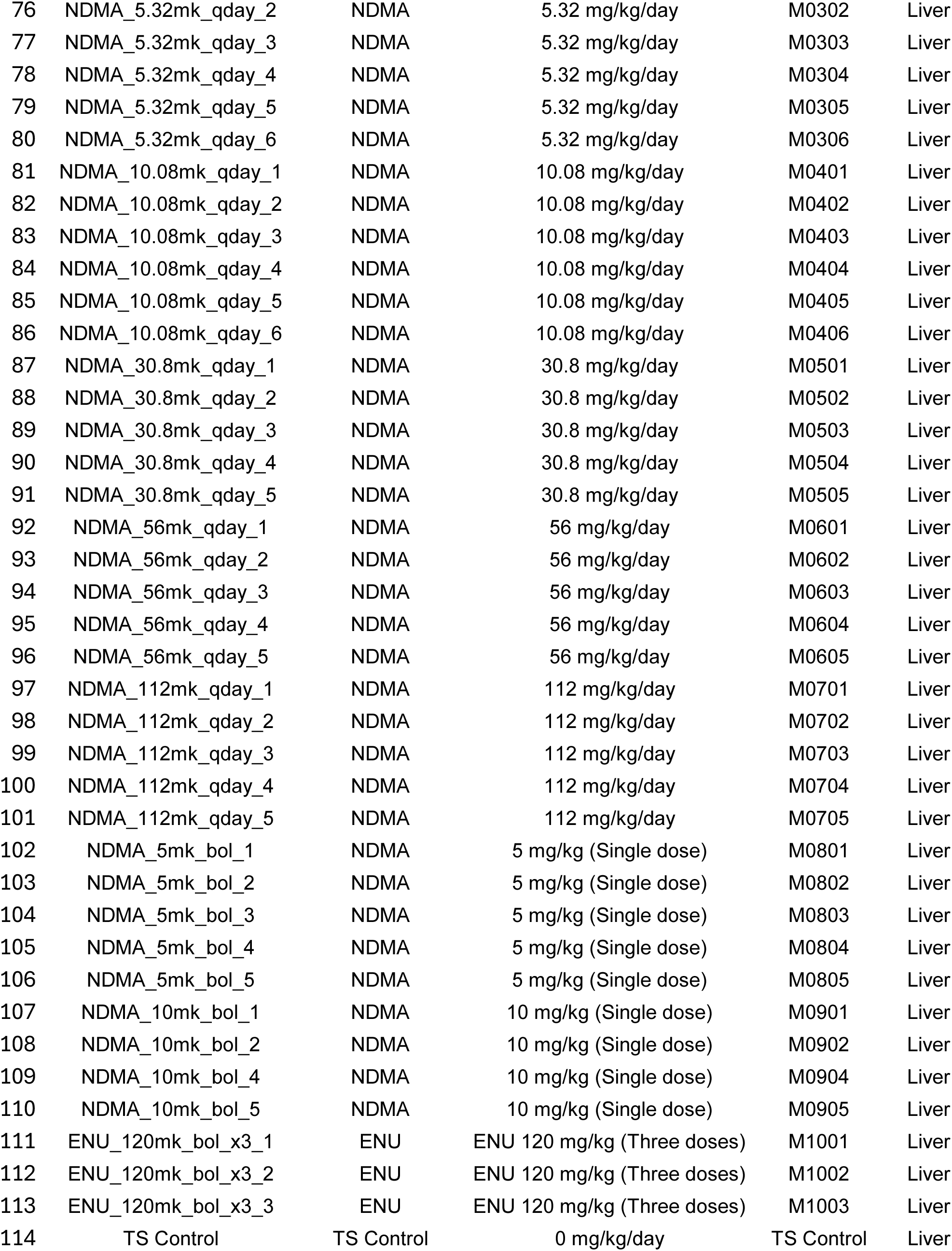

| Row | Sample_Alias | Treatment | Dose | Animal | Tissue |
| --- | --- | --- | --- | --- | --- |
| 1 | Vehicle Control_1 | Vehicle Control | 0 mg/kg/day | M0001 | Liver |
| 2 | Vehicle Control_2 | Vehicle Control | 0 mg/kg/day | M0002 | Liver |
| 3 | Vehicle Control_3 | Vehicle Control | 0 mg/kg/day | M0003 | Liver |
| 4 | Vehicle Control_4 | Vehicle Control | 0 mg/kg/day | M0004 | Liver |
| 5 | Vehicle Control_5 | Vehicle Control | 0 mg/kg/day | M0005 | Liver |
| 6 | Vehicle Control_6 | Vehicle Control | 0 mg/kg/day | M0006 | Liver |
| 7 | NDMA_0.56mk_qday_1 | NDMA | 0.56 mg/kg/day | M0101 | Liver |
| 8 | NDMA_0.56mk_qday_2 | NDMA | 0.56 mg/kg/day | M0102 | Liver |
| 9 | NDMA_0.56mk_qday_3 | NDMA | 0.56 mg/kg/day | M0103 | Liver |
| 10 | NDMA_0.56mk_qday_4 | NDMA | 0.56 mg/kg/day | M0104 | Liver |
| 11 | NDMA_0.56mk_qday_5 | NDMA | 0.56 mg/kg/day | M0105 | Liver |
| 12 | NDMA_1.96mk_qday_1 | NDMA | 1.96 mg/kg/day | M0201 | Liver |
| 13 | NDMA_1.96mk_qday_2 | NDMA | 1.96 mg/kg/day | M0202 | Liver |
| 14 | NDMA_1.96mk_qday_3 | NDMA | 1.96 mg/kg/day | M0203 | Liver |
| 15 | NDMA_1.96mk_qday_4 | NDMA | 1.96 mg/kg/day | M0204 | Liver |
| 16 | NDMA_1.96mk_qday_5 | NDMA | 1.96 mg/kg/day | M0205 | Liver |
| 17 | NDMA_1.96mk_qday_6 | NDMA | 1.96 mg/kg/day | M0206 | Liver |
| 18 | NDMA_5.32mk_qday_1 | NDMA | 5.32 mg/kg/day | M0301 | Liver |
| 19 | NDMA_5.32mk_qday_2 | NDMA | 5.32 mg/kg/day | M0302 | Liver |
| 20 | NDMA_5.32mk_qday_3 | NDMA | 5.32 mg/kg/day | M0303 | Liver |
| 21 | NDMA_5.32mk_qday_4 | NDMA | 5.32 mg/kg/day | M0304 | Liver |
| 22 | NDMA_5.32mk_qday_5 | NDMA | 5.32 mg/kg/day | M0305 | Liver |
| 23 | NDMA_5.32mk_qday_6 | NDMA | 5.32 mg/kg/day | M0306 | Liver |
| 24 | NDMA_10.08mk_qday_1 | NDMA | 10.08 mg/kg/day | M0401 | Liver |
| 25 | NDMA_10.08mk_qday_2 | NDMA | 10.08 mg/kg/day | M0402 | Liver |
| 26 | NDMA_10.08mk_qday_3 | NDMA | 10.08 mg/kg/day | M0403 | Liver |
| 27 | NDMA_10.08mk_qday_4 | NDMA | 10.08 mg/kg/day | M0404 | Liver |
| 28 | NDMA_10.08mk_qday_5 | NDMA | 10.08 mg/kg/day | M0405 | Liver |
| 29 | NDMA_10.08mk_qday_6 | NDMA | 10.08 mg/kg/day | M0406 | Liver |
| 30 | NDMA_30.8mk_qday_1 | NDMA | 30.8 mg/kg/day | M0501 | Liver |
| 31 | NDMA_30.8mk_qday_2 | NDMA | 30.8 mg/kg/day | M0502 | Liver |
| 32 | NDMA_30.8mk_qday_3 | NDMA | 30.8 mg/kg/day | M0503 | Liver |
| 33 | NDMA_30.8mk_qday_4 | NDMA | 30.8 mg/kg/day | M0504 | Liver |
| 34 | NDMA_30.8mk_qday_5 | NDMA | 30.8 mg/kg/day | M0505 | Liver |
| 35 | NDMA_56mk_qday_1 | NDMA | 56 mg/kg/day | M0601 | Liver |
| 36 | NDMA_56mk_qday_2 | NDMA | 56 mg/kg/day | M0602 | Liver |
| 37 | NDMA_56mk_qday_3 | NDMA | 56 mg/kg/day | M0603 | Liver |
| 38 | NDMA_56mk_qday_4 | NDMA | 56 mg/kg/day | M0604 | Liver |
| 39 | NDMA_56mk_qday_5 | NDMA | 56 mg/kg/day | M0605 | Liver |
| 40 | NDMA_112mk_qday_1 | NDMA | 112 mg/kg/day | M0701 | Liver |
| 41 | NDMA_112mk_qday_2 | NDMA | 112 mg/kg/day | M0702 | Liver |
| 42 | NDMA_112mk_qday_3 | NDMA | 112 mg/kg/day | M0703 | Liver |
| 43 | NDMA_112mk_qday_4 | NDMA | 112 mg/kg/day | M0704 | Liver |
| 44 | NDMA_112mk_qday_5 | NDMA | 112 mg/kg/day | M0705 | Liver |
| 45 | NDMA_5mk_bol_1 | NDMA | 5 mg/kg (Single dose) | M0801 | Liver |
| 46 | NDMA_5mk_bol_2 | NDMA | 5 mg/kg (Single dose) | M0802 | Liver |
| 47 | NDMA_5mk_bol_3 | NDMA | 5 mg/kg (Single dose) | M0803 | Liver |
| 48 | NDMA_5mk_bol_4 | NDMA | 5 mg/kg (Single dose) | M0804 | Liver |
| 49 | NDMA_5mk_bol_5 | NDMA | 5 mg/kg (Single dose) | M0805 | Liver |
| 50 | NDMA_10mk_bol_1 | NDMA | 10 mg/kg (Single dose) | M0901 | Liver |
| 51 | NDMA_10mk_bol_2 | NDMA | 10 mg/kg (Single dose) | M0902 | Liver |
| 52 | NDMA_10mk_bol_4 | NDMA | 10 mg/kg (Single dose) | M0904 | Liver |
| 53 | NDMA_10mk_bol_5 | NDMA | 10 mg/kg (Single dose) | M0905 | Liver |
| 54 | ENU_120mk_bol_x3_1 | ENU | ENU 120 mg/kg (Three doses) | M1001 | Liver |
| 55 | ENU_120mk_bol_x3_2 | ENU | ENU 120 mg/kg (Three doses) | M1002 | Liver |
| 56 | ENU_120mk_bol_x3_3 | ENU | ENU 120 mg/kg (Three doses) | M1003 | Liver |
| 57 | TS Control | TS Control | 0 mg/kg/day | TS Control | Liver |
| 58 | Vehicle Control_1 | Vehicle Control | 0 mg/kg/day | M0001 | Liver |
| 59 | Vehicle Control_2 | Vehicle Control | 0 mg/kg/day | M0002 | Liver |
| 60 | Vehicle Control_3 | Vehicle Control | 0 mg/kg/day | M0003 | Liver |
| 61 | Vehicle Control_4 | Vehicle Control | 0 mg/kg/day | M0004 | Liver |
| 62 | Vehicle Control_5 | Vehicle Control | 0 mg/kg/day | M0005 | Liver |
| 63 | Vehicle Control_6 | Vehicle Control | 0 mg/kg/day | M0006 | Liver |
| 64 | NDMA_0.56mk_qday_1 | NDMA | 0.56 mg/kg/day | M0101 | Liver |
| 65 | NDMA_0.56mk_qday_2 | NDMA | 0.56 mg/kg/day | M0102 | Liver |
| 66 | NDMA_0.56mk_qday_3 | NDMA | 0.56 mg/kg/day | M0103 | Liver |
| 67 | NDMA_0.56mk_qday_4 | NDMA | 0.56 mg/kg/day | M0104 | Liver |
| 68 | NDMA_0.56mk_qday_5 | NDMA | 0.56 mg/kg/day | M0105 | Liver |
| 69 | NDMA_1.96mk_qday_1 | NDMA | 1.96 mg/kg/day | M0201 | Liver |
| 70 | NDMA_1.96mk_qday_2 | NDMA | 1.96 mg/kg/day | M0202 | Liver |
| 71 | NDMA_1.96mk_qday_3 | NDMA | 1.96 mg/kg/day | M0203 | Liver |
| 72 | NDMA_1.96mk_qday_4 | NDMA | 1.96 mg/kg/day | M0204 | Liver |
| 73 | NDMA_1.96mk_qday_5 | NDMA | 1.96 mg/kg/day | M0205 | Liver |
| 74 | NDMA_1.96mk_qday_6 | NDMA | 1.96 mg/kg/day | M0206 | Liver |
| 75 | NDMA_5.32mk_qday_1 | NDMA | 5.32 mg/kg/day | M0301 | Liver |
| 76 | NDMA_5.32mk_qday_2 | NDMA | 5.32 mg/kg/day | M0302 | Liver |
| 77 | NDMA_5.32mk_qday_3 | NDMA | 5.32 mg/kg/day | M0303 | Liver |
| 78 | NDMA_5.32mk_qday_4 | NDMA | 5.32 mg/kg/day | M0304 | Liver |
| 79 | NDMA_5.32mk_qday_5 | NDMA | 5.32 mg/kg/day | M0305 | Liver |
| 80 | NDMA_5.32mk_qday_6 | NDMA | 5.32 mg/kg/day | M0306 | Liver |
| 81 | NDMA_10.08mk_qday_1 | NDMA | 10.08 mg/kg/day | M0401 | Liver |
| 82 | NDMA_10.08mk_qday_2 | NDMA | 10.08 mg/kg/day | M0402 | Liver |
| 83 | NDMA_10.08mk_qday_3 | NDMA | 10.08 mg/kg/day | M0403 | Liver |
| 84 | NDMA_10.08mk_qday_4 | NDMA | 10.08 mg/kg/day | M0404 | Liver |
| 85 | NDMA_10.08mk_qday_5 | NDMA | 10.08 mg/kg/day | M0405 | Liver |
| 86 | NDMA_10.08mk_qday_6 | NDMA | 10.08 mg/kg/day | M0406 | Liver |
| 87 | NDMA_30.8mk_qday_1 | NDMA | 30.8 mg/kg/day | M0501 | Liver |
| 88 | NDMA_30.8mk_qday_2 | NDMA | 30.8 mg/kg/day | M0502 | Liver |
| 89 | NDMA_30.8mk_qday_3 | NDMA | 30.8 mg/kg/day | M0503 | Liver |
| 90 | NDMA_30.8mk_qday_4 | NDMA | 30.8 mg/kg/day | M0504 | Liver |
| 91 | NDMA_30.8mk_qday_5 | NDMA | 30.8 mg/kg/day | M0505 | Liver |
| 92 | NDMA_56mk_qday_1 | NDMA | 56 mg/kg/day | M0601 | Liver |
| 93 | NDMA_56mk_qday_2 | NDMA | 56 mg/kg/day | M0602 | Liver |
| 94 | NDMA_56mk_qday_3 | NDMA | 56 mg/kg/day | M0603 | Liver |
| 95 | NDMA_56mk_qday_4 | NDMA | 56 mg/kg/day | M0604 | Liver |
| 96 | NDMA_56mk_qday_5 | NDMA | 56 mg/kg/day | M0605 | Liver |
| 97 | NDMA_112mk_qday_1 | NDMA | 112 mg/kg/day | M0701 | Liver |
| 98 | NDMA_112mk_qday_2 | NDMA | 112 mg/kg/day | M0702 | Liver |
| 99 | NDMA_112mk_qday_3 | NDMA | 112 mg/kg/day | M0703 | Liver |
| 100 | NDMA_112mk_qday_4 | NDMA | 112 mg/kg/day | M0704 | Liver |
| 101 | NDMA_112mk_qday_5 | NDMA | 112 mg/kg/day | M0705 | Liver |
| 102 | NDMA_5mk_bol_1 | NDMA | 5 mg/kg (Single dose) | M0801 | Liver |
| 103 | NDMA_5mk_bol_2 | NDMA | 5 mg/kg (Single dose) | M0802 | Liver |
| 104 | NDMA_5mk_bol_3 | NDMA | 5 mg/kg (Single dose) | M0803 | Liver |
| 105 | NDMA_5mk_bol_4 | NDMA | 5 mg/kg (Single dose) | M0804 | Liver |
| 106 | NDMA_5mk_bol_5 | NDMA | 5 mg/kg (Single dose) | M0805 | Liver |
| 107 | NDMA_10mk_bol_1 | NDMA | 10 mg/kg (Single dose) | M0901 | Liver |
| 108 | NDMA_10mk_bol_2 | NDMA | 10 mg/kg (Single dose) | M0902 | Liver |
| 109 | NDMA_10mk_bol_4 | NDMA | 10 mg/kg (Single dose) | M0904 | Liver |
| 110 | NDMA_10mk_bol_5 | NDMA | 10 mg/kg (Single dose) | M0905 | Liver |
| 111 | ENU_120mk_bol_x3_1 | ENU | ENU 120 mg/kg (Three doses) | M1001 | Liver |
| 112 | ENU_120mk_bol_x3_2 | ENU | ENU 120 mg/kg (Three doses) | M1002 | Liver |
| 113 | ENU_120mk_bol_x3_3 | ENU | ENU 120 mg/kg (Three doses) | M1003 | Liver |
| 114 | TS Control | TS Control | 0 mg/kg/day | TS Control | Liver |

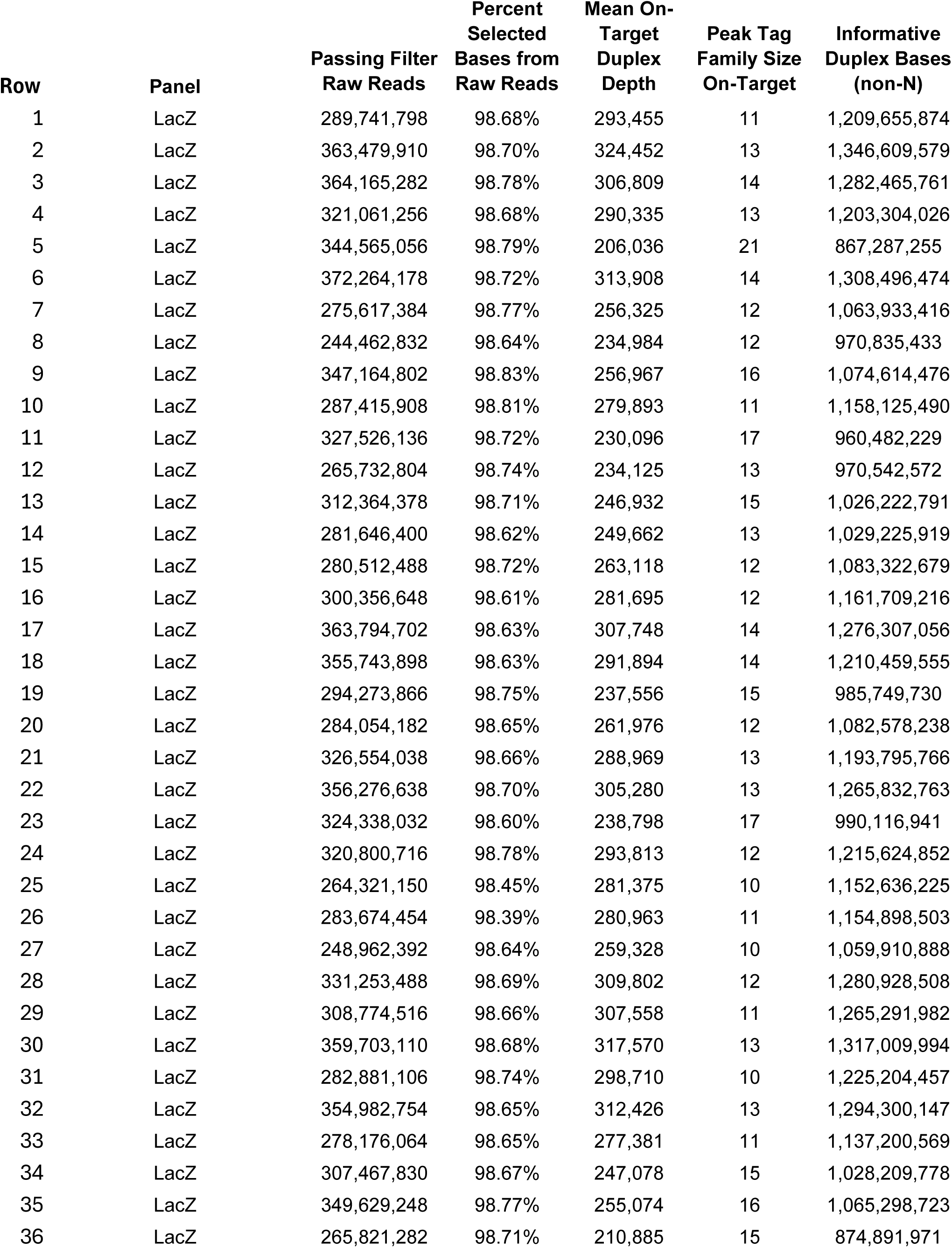
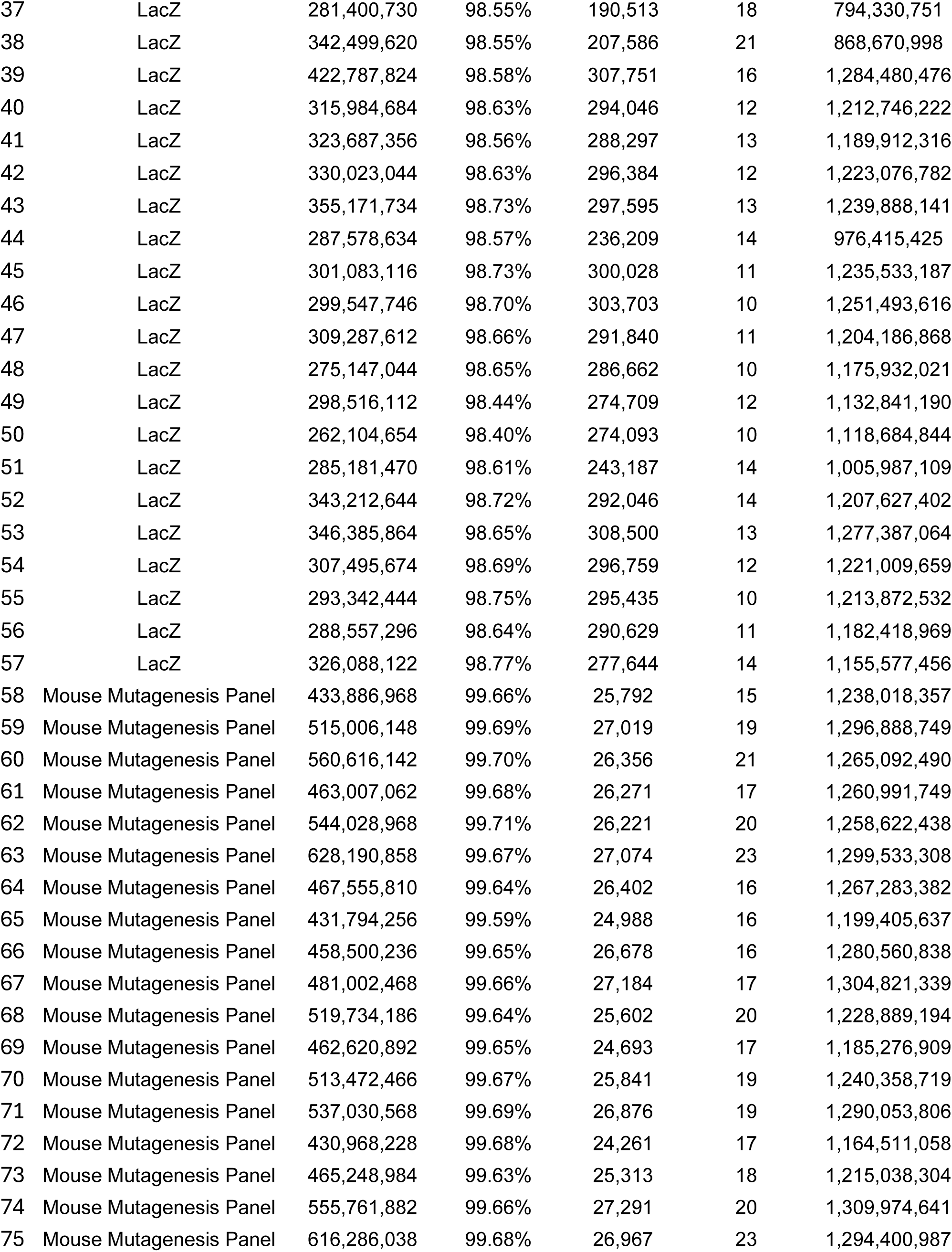
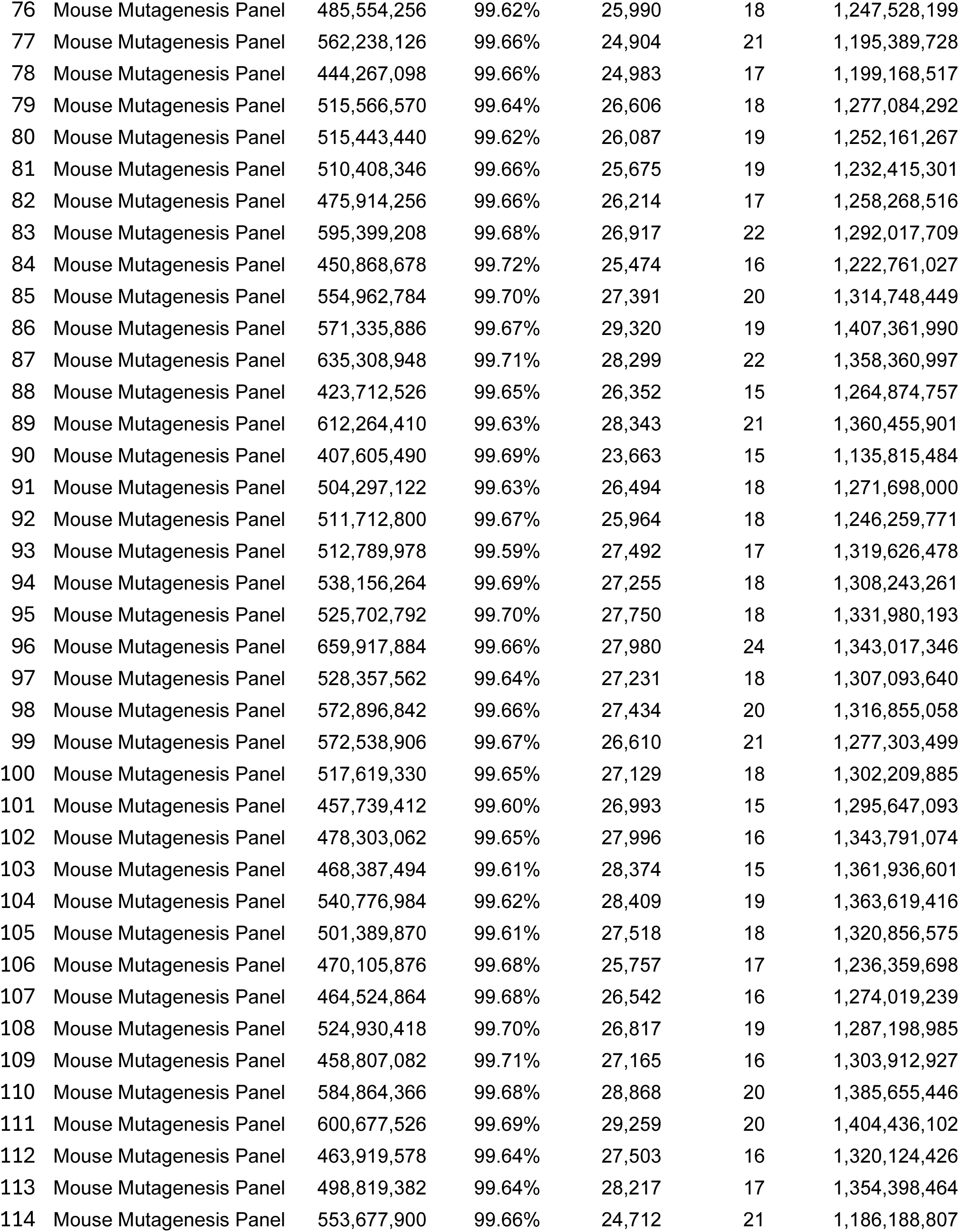
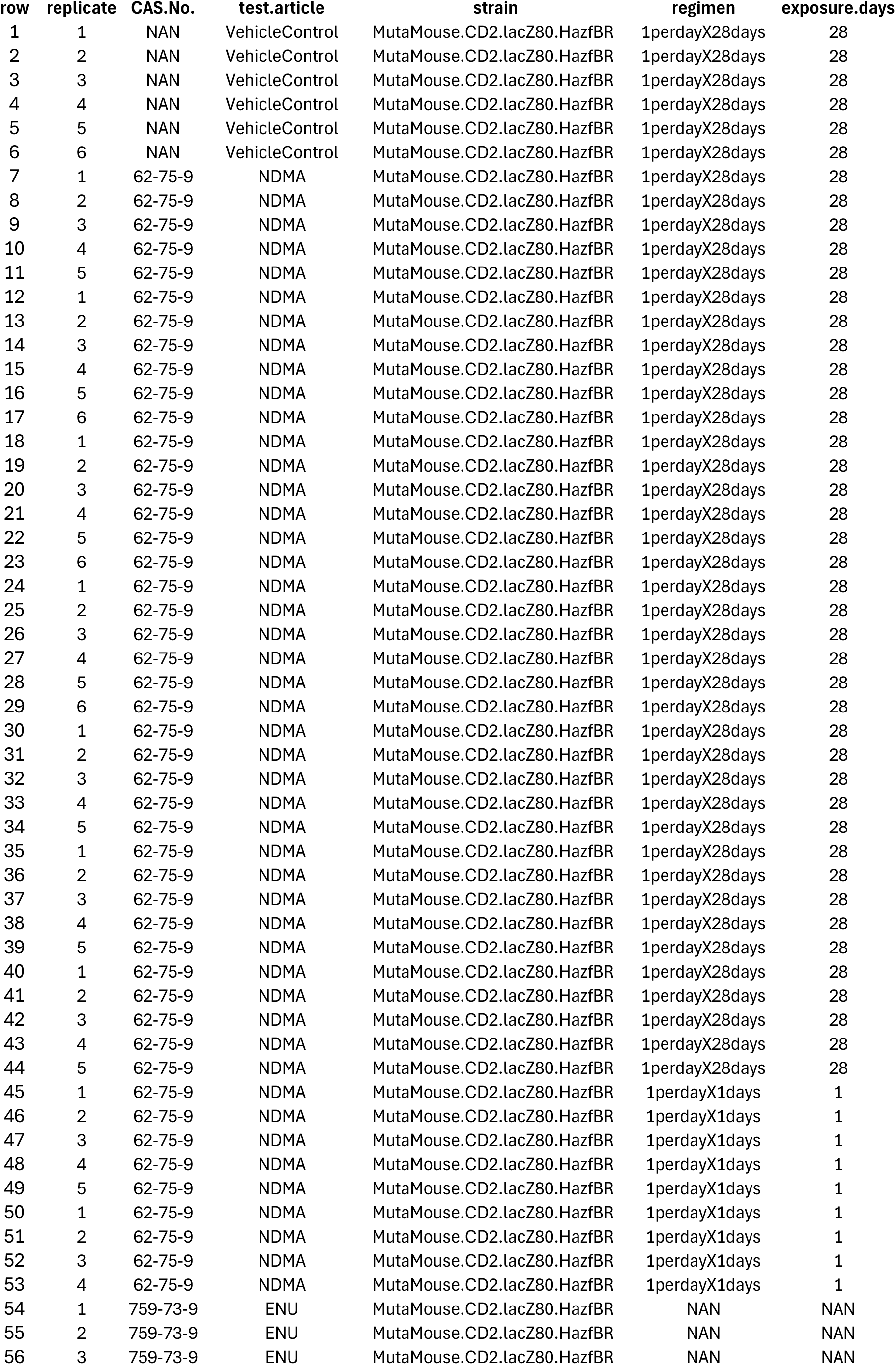
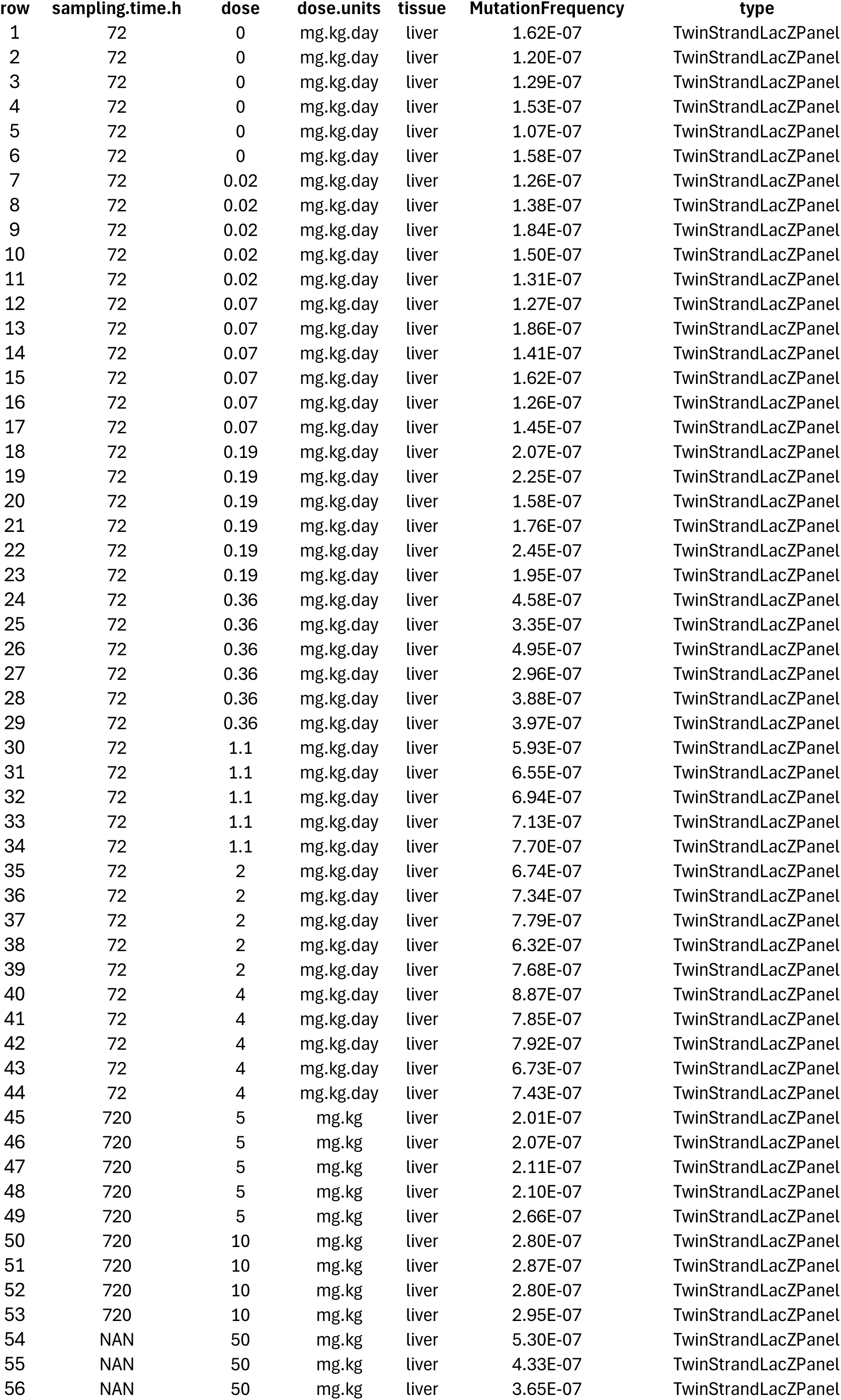
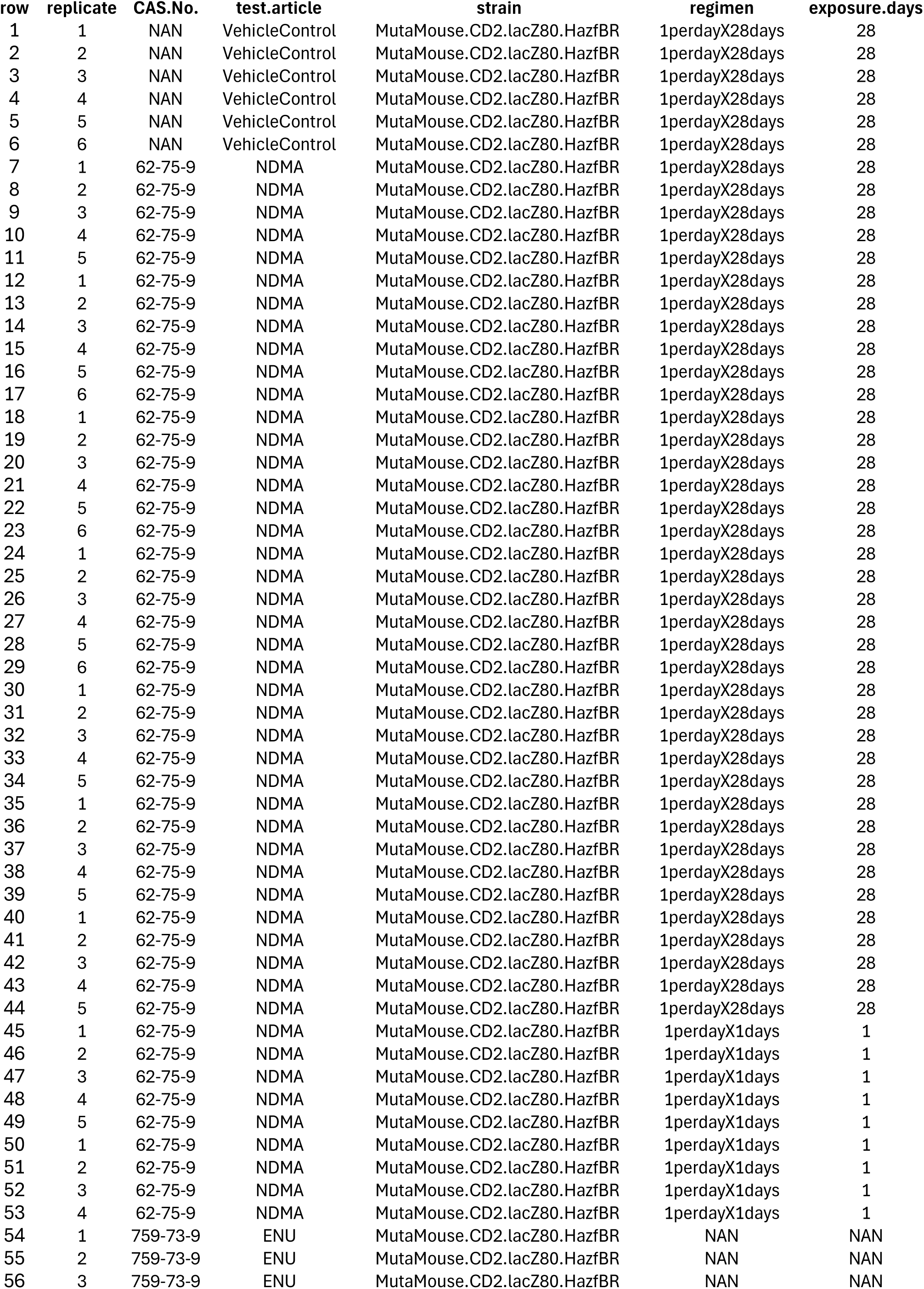
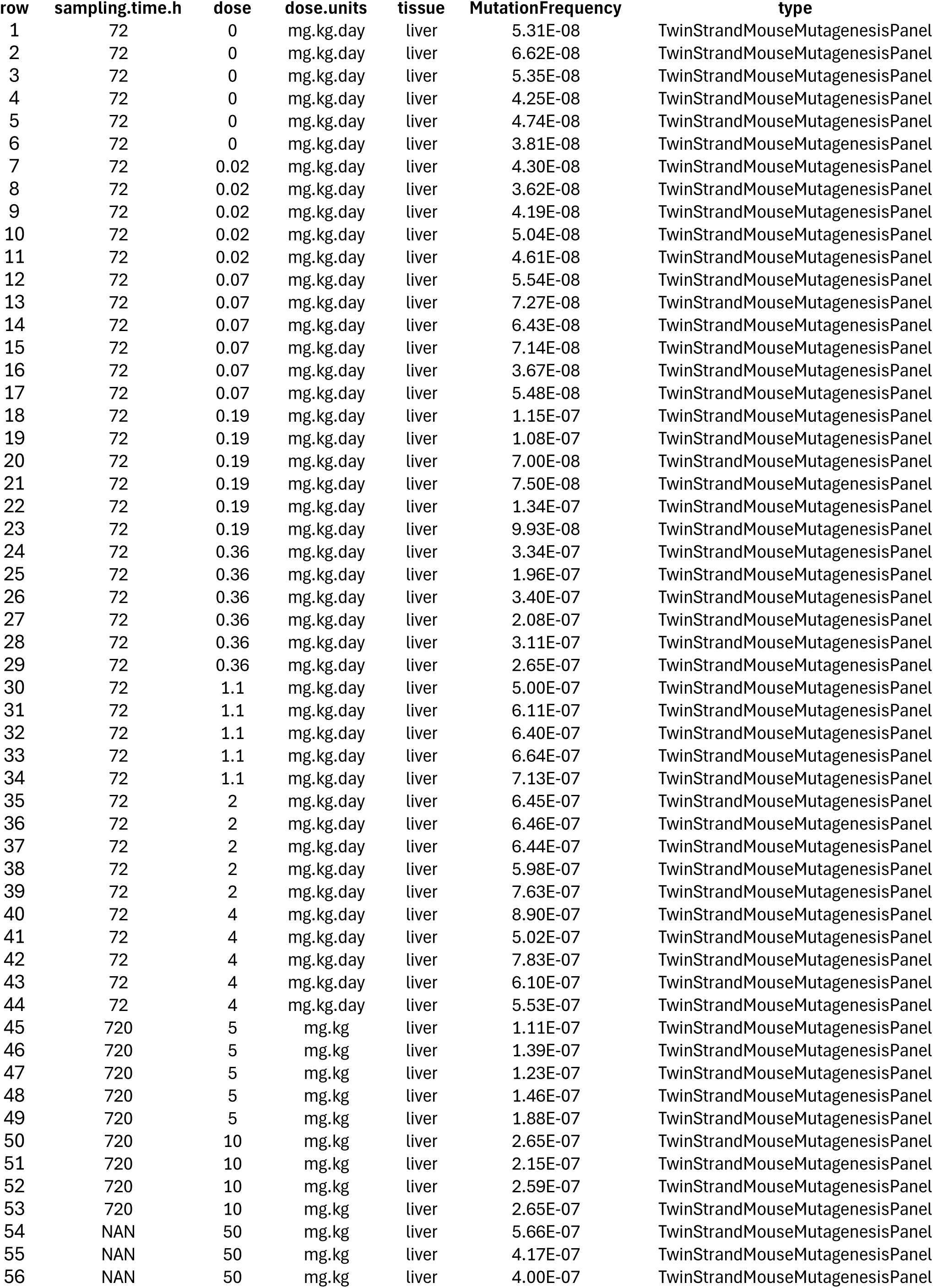

| Row | Panel | Passing Filter<br>Raw Reads | Percent<br>Selected<br>Bases from<br>Raw Reads | Mean On-Target<br>Duplex<br>Depth | Peak Tag<br>Family Size<br>On-Target | Informative<br>Duplex Bases<br>(non-N) |
| --- | --- | --- | --- | --- | --- | --- |
| 1 | LacZ | 289,741,798 | 98.68% | 293,455 | 11 | 1,209,655,874 |
| 2 | LacZ | 363,479,910 | 98.70% | 324,452 | 13 | 1,346,609,579 |
| 3 | LacZ | 364,165,282 | 98.78% | 306,809 | 14 | 1,282,465,761 |
| 4 | LacZ | 321,061,256 | 98.68% | 290,335 | 13 | 1,203,304,026 |
| 5 | LacZ | 344,565,056 | 98.79% | 206,036 | 21 | 867,287,255 |
| 6 | LacZ | 372,264,178 | 98.72% | 313,908 | 14 | 1,308,496,474 |
| 7 | LacZ | 275,617,384 | 98.77% | 256,325 | 12 | 1,063,933,416 |
| 8 | LacZ | 244,462,832 | 98.64% | 234,984 | 12 | 970,835,433 |
| 9 | LacZ | 347,164,802 | 98.83% | 256,967 | 16 | 1,074,614,476 |
| 10 | LacZ | 287,415,908 | 98.81% | 279,893 | 11 | 1,158,125,490 |
| 11 | LacZ | 327,526,136 | 98.72% | 230,096 | 17 | 960,482,229 |
| 12 | LacZ | 265,732,804 | 98.74% | 234,125 | 13 | 970,542,572 |
| 13 | LacZ | 312,364,378 | 98.71% | 246,932 | 15 | 1,026,222,791 |
| 14 | LacZ | 281,646,400 | 98.62% | 249,662 | 13 | 1,029,225,919 |
| 15 | LacZ | 280,512,488 | 98.72% | 263,118 | 12 | 1,083,322,679 |
| 16 | LacZ | 300,356,648 | 98.61% | 281,695 | 12 | 1,161,709,216 |
| 17 | LacZ | 363,794,702 | 98.63% | 307,748 | 14 | 1,276,307,056 |
| 18 | LacZ | 355,743,898 | 98.63% | 291,894 | 14 | 1,210,459,555 |
| 19 | LacZ | 294,273,866 | 98.75% | 237,556 | 15 | 985,749,730 |
| 20 | LacZ | 284,054,182 | 98.65% | 261,976 | 12 | 1,082,578,238 |
| 21 | LacZ | 326,554,038 | 98.66% | 288,969 | 13 | 1,193,795,766 |
| 22 | LacZ | 356,276,638 | 98.70% | 305,280 | 13 | 1,265,832,763 |
| 23 | LacZ | 324,338,032 | 98.60% | 238,798 | 17 | 990,116,941 |
| 24 | LacZ | 320,800,716 | 98.78% | 293,813 | 12 | 1,215,624,852 |
| 25 | LacZ | 264,321,150 | 98.45% | 281,375 | 10 | 1,152,636,225 |
| 26 | LacZ | 283,674,454 | 98.39% | 280,963 | 11 | 1,154,898,503 |
| 27 | LacZ | 248,962,392 | 98.64% | 259,328 | 10 | 1,059,910,888 |
| 28 | LacZ | 331,253,488 | 98.69% | 309,802 | 12 | 1,280,928,508 |
| 29 | LacZ | 308,774,516 | 98.66% | 307,558 | 11 | 1,265,291,982 |
| 30 | LacZ | 359,703,110 | 98.68% | 317,570 | 13 | 1,317,009,994 |
| 31 | LacZ | 282,881,106 | 98.74% | 298,710 | 10 | 1,225,204,457 |
| 32 | LacZ | 354,982,754 | 98.65% | 312,426 | 13 | 1,294,300,147 |
| 33 | LacZ | 278,176,064 | 98.65% | 277,381 | 11 | 1,137,200,569 |
| 34 | LacZ | 307,467,830 | 98.67% | 247,078 | 15 | 1,028,209,778 |
| 35 | LacZ | 349,629,248 | 98.77% | 255,074 | 16 | 1,065,298,723 |
| 36 | LacZ | 265,821,282 | 98.71% | 210,885 | 15 | 874,891,971 |
| 37 | LacZ | 281,400,730 | 98.55% | 190,513 | 18 | 794,330,751 |
| 38 | LacZ | 342,499,620 | 98.55% | 207,586 | 21 | 868,670,998 |
| 39 | LacZ | 422,787,824 | 98.58% | 307,751 | 16 | 1,284,480,476 |
| 40 | LacZ | 315,984,684 | 98.63% | 294,046 | 12 | 1,212,746,222 |
| 41 | LacZ | 323,687,356 | 98.56% | 288,297 | 13 | 1,189,912,316 |
| 42 | LacZ | 330,023,044 | 98.63% | 296,384 | 12 | 1,223,076,782 |
| 43 | LacZ | 355,171,734 | 98.73% | 297,595 | 13 | 1,239,888,141 |
| 44 | LacZ | 287,578,634 | 98.57% | 236,209 | 14 | 976,415,425 |
| 45 | LacZ | 301,083,116 | 98.73% | 300,028 | 11 | 1,235,533,187 |
| 46 | LacZ | 299,547,746 | 98.70% | 303,703 | 10 | 1,251,493,616 |
| 47 | LacZ | 309,287,612 | 98.66% | 291,840 | 11 | 1,204,186,868 |
| 48 | LacZ | 275,147,044 | 98.65% | 286,662 | 10 | 1,175,932,021 |
| 49 | LacZ | 298,516,112 | 98.44% | 274,709 | 12 | 1,132,841,190 |
| 50 | LacZ | 262,104,654 | 98.40% | 274,093 | 10 | 1,118,684,844 |
| 51 | LacZ | 285,181,470 | 98.61% | 243,187 | 14 | 1,005,987,109 |
| 52 | LacZ | 343,212,644 | 98.72% | 292,046 | 14 | 1,207,627,402 |
| 53 | LacZ | 346,385,864 | 98.65% | 308,500 | 13 | 1,277,387,064 |
| 54 | LacZ | 307,495,674 | 98.69% | 296,759 | 12 | 1,221,009,659 |
| 55 | LacZ | 293,342,444 | 98.75% | 295,435 | 10 | 1,213,872,532 |
| 56 | LacZ | 288,557,296 | 98.64% | 290,629 | 11 | 1,182,418,969 |
| 57 | LacZ | 326,088,122 | 98.77% | 277,644 | 14 | 1,155,577,456 |
| 58 | Mouse Mutagenesis Panel | 433,886,968 | 99.66% | 25,792 | 15 | 1,238,018,357 |
| 59 | Mouse Mutagenesis Panel | 515,006,148 | 99.69% | 27,019 | 19 | 1,296,888,749 |
| 60 | Mouse Mutagenesis Panel | 560,616,142 | 99.70% | 26,356 | 21 | 1,265,092,490 |
| 61 | Mouse Mutagenesis Panel | 463,007,062 | 99.68% | 26,271 | 17 | 1,260,991,749 |
| 62 | Mouse Mutagenesis Panel | 544,028,968 | 99.71% | 26,221 | 20 | 1,258,622,438 |
| 63 | Mouse Mutagenesis Panel | 628,190,858 | 99.67% | 27,074 | 23 | 1,299,533,308 |
| 64 | Mouse Mutagenesis Panel | 467,555,810 | 99.64% | 26,402 | 16 | 1,267,283,382 |
| 65 | Mouse Mutagenesis Panel | 431,794,256 | 99.59% | 24,988 | 16 | 1,199,405,637 |
| 66 | Mouse Mutagenesis Panel | 458,500,236 | 99.65% | 26,678 | 16 | 1,280,560,838 |
| 67 | Mouse Mutagenesis Panel | 481,002,468 | 99.66% | 27,184 | 17 | 1,304,821,339 |
| 68 | Mouse Mutagenesis Panel | 519,734,186 | 99.64% | 25,602 | 20 | 1,228,889,194 |
| 69 | Mouse Mutagenesis Panel | 462,620,892 | 99.65% | 24,693 | 17 | 1,185,276,909 |
| 70 | Mouse Mutagenesis Panel | 513,472,466 | 99.67% | 25,841 | 19 | 1,240,358,719 |
| 71 | Mouse Mutagenesis Panel | 537,030,568 | 99.69% | 26,876 | 19 | 1,290,053,806 |
| 72 | Mouse Mutagenesis Panel | 430,968,228 | 99.68% | 24,261 | 17 | 1,164,511,058 |
| 73 | Mouse Mutagenesis Panel | 465,248,984 | 99.63% | 25,313 | 18 | 1,215,038,304 |
| 74 | Mouse Mutagenesis Panel | 555,761,882 | 99.66% | 27,291 | 20 | 1,309,974,641 |
| 75 | Mouse Mutagenesis Panel | 616,286,038 | 99.68% | 26,967 | 23 | 1,294,400,987 |
| 76 | Mouse Mutagenesis Panel | 485,554,256 | 99.62% | 25,990 | 18 | 1,247,528,199 |
| 77 | Mouse Mutagenesis Panel | 562,238,126 | 99.66% | 24,904 | 21 | 1,195,389,728 |
| 78 | Mouse Mutagenesis Panel | 444,267,098 | 99.66% | 24,983 | 17 | 1,199,168,517 |
| 79 | Mouse Mutagenesis Panel | 515,566,570 | 99.64% | 26,606 | 18 | 1,277,084,292 |
| 80 | Mouse Mutagenesis Panel | 515,443,440 | 99.62% | 26,087 | 19 | 1,252,161,267 |
| 81 | Mouse Mutagenesis Panel | 510,408,346 | 99.66% | 25,675 | 19 | 1,232,415,301 |
| 82 | Mouse Mutagenesis Panel | 475,914,256 | 99.66% | 26,214 | 17 | 1,258,268,516 |
| 83 | Mouse Mutagenesis Panel | 595,399,208 | 99.68% | 26,917 | 22 | 1,292,017,709 |
| 84 | Mouse Mutagenesis Panel | 450,868,678 | 99.72% | 25,474 | 16 | 1,222,761,027 |
| 85 | Mouse Mutagenesis Panel | 554,962,784 | 99.70% | 27,391 | 20 | 1,314,748,449 |
| 86 | Mouse Mutagenesis Panel | 571,335,886 | 99.67% | 29,320 | 19 | 1,407,361,990 |
| 87 | Mouse Mutagenesis Panel | 635,308,948 | 99.71% | 28,299 | 22 | 1,358,360,997 |
| 88 | Mouse Mutagenesis Panel | 423,712,526 | 99.65% | 26,352 | 15 | 1,264,874,757 |
| 89 | Mouse Mutagenesis Panel | 612,264,410 | 99.63% | 28,343 | 21 | 1,360,455,901 |
| 90 | Mouse Mutagenesis Panel | 407,605,490 | 99.69% | 23,663 | 15 | 1,135,815,484 |
| 91 | Mouse Mutagenesis Panel | 504,297,122 | 99.63% | 26,494 | 18 | 1,271,698,000 |
| 92 | Mouse Mutagenesis Panel | 511,712,800 | 99.67% | 25,964 | 18 | 1,246,259,771 |
| 93 | Mouse Mutagenesis Panel | 512,789,978 | 99.59% | 27,492 | 17 | 1,319,626,478 |
| 94 | Mouse Mutagenesis Panel | 538,156,264 | 99.69% | 27,255 | 18 | 1,308,243,261 |
| 95 | Mouse Mutagenesis Panel | 525,702,792 | 99.70% | 27,750 | 18 | 1,331,980,193 |
| 96 | Mouse Mutagenesis Panel | 659,917,884 | 99.66% | 27,980 | 24 | 1,343,017,346 |
| 97 | Mouse Mutagenesis Panel | 528,357,562 | 99.64% | 27,231 | 18 | 1,307,093,640 |
| 98 | Mouse Mutagenesis Panel | 572,896,842 | 99.66% | 27,434 | 20 | 1,316,855,058 |
| 99 | Mouse Mutagenesis Panel | 572,538,906 | 99.67% | 26,610 | 21 | 1,277,303,499 |
| 100 | Mouse Mutagenesis Panel | 517,619,330 | 99.65% | 27,129 | 18 | 1,302,209,885 |
| 101 | Mouse Mutagenesis Panel | 457,739,412 | 99.60% | 26,993 | 15 | 1,295,647,093 |
| 102 | Mouse Mutagenesis Panel | 478,303,062 | 99.65% | 27,996 | 16 | 1,343,791,074 |
| 103 | Mouse Mutagenesis Panel | 468,387,494 | 99.61% | 28,374 | 15 | 1,361,936,601 |
| 104 | Mouse Mutagenesis Panel | 540,776,984 | 99.62% | 28,409 | 19 | 1,363,619,416 |
| 105 | Mouse Mutagenesis Panel | 501,389,870 | 99.61% | 27,518 | 18 | 1,320,856,575 |
| 106 | Mouse Mutagenesis Panel | 470,105,876 | 99.68% | 25,757 | 17 | 1,236,359,698 |
| 107 | Mouse Mutagenesis Panel | 464,524,864 | 99.68% | 26,542 | 16 | 1,274,019,239 |
| 108 | Mouse Mutagenesis Panel | 524,930,418 | 99.70% | 26,817 | 19 | 1,287,198,985 |
| 109 | Mouse Mutagenesis Panel | 458,807,082 | 99.71% | 27,165 | 16 | 1,303,912,927 |
| 110 | Mouse Mutagenesis Panel | 584,864,366 | 99.68% | 28,868 | 20 | 1,385,655,446 |
| 111 | Mouse Mutagenesis Panel | 600,677,526 | 99.69% | 29,259 | 20 | 1,404,436,102 |
| 112 | Mouse Mutagenesis Panel | 463,919,578 | 99.64% | 27,503 | 16 | 1,320,124,426 |
| 113 | Mouse Mutagenesis Panel | 498,819,382 | 99.64% | 28,217 | 17 | 1,354,398,464 |
| 114 | Mouse Mutagenesis Panel | 553,677,900 | 99.66% | 24,712 | 21 | 1,186,188,807 |

| row | replicate | CAS.No. | test.article | strain | regimen | exposure.days |
| --- | --- | --- | --- | --- | --- | --- |
| 1 | 1 | NAN | VehicleControl | MutaMouse.CD2.lacZ80.HazfBR | 1perdayX28days | 28 |
| 2 | 2 | NAN | VehicleControl | MutaMouse.CD2.lacZ80.HazfBR | 1perdayX28days | 28 |
| 3 | 3 | NAN | VehicleControl | MutaMouse.CD2.lacZ80.HazfBR | 1perdayX28days | 28 |
| 4 | 4 | NAN | VehicleControl | MutaMouse.CD2.lacZ80.HazfBR | 1perdayX28days | 28 |
| 5 | 5 | NAN | VehicleControl | MutaMouse.CD2.lacZ80.HazfBR | 1perdayX28days | 28 |
| 6 | 6 | NAN | VehicleControl | MutaMouse.CD2.lacZ80.HazfBR | 1perdayX28days | 28 |
| 7 | 1 | 62-75-9 | NDMA | MutaMouse.CD2.lacZ80.HazfBR | 1perdayX28days | 28 |
| 8 | 2 | 62-75-9 | NDMA | MutaMouse.CD2.lacZ80.HazfBR | 1perdayX28days | 28 |
| 9 | 3 | 62-75-9 | NDMA | MutaMouse.CD2.lacZ80.HazfBR | 1perdayX28days | 28 |
| 10 | 4 | 62-75-9 | NDMA | MutaMouse.CD2.lacZ80.HazfBR | 1perdayX28days | 28 |
| 11 | 5 | 62-75-9 | NDMA | MutaMouse.CD2.lacZ80.HazfBR | 1perdayX28days | 28 |
| 12 | 1 | 62-75-9 | NDMA | MutaMouse.CD2.lacZ80.HazfBR | 1perdayX28days | 28 |
| 13 | 2 | 62-75-9 | NDMA | MutaMouse.CD2.lacZ80.HazfBR | 1perdayX28days | 28 |
| 14 | 3 | 62-75-9 | NDMA | MutaMouse.CD2.lacZ80.HazfBR | 1perdayX28days | 28 |
| 15 | 4 | 62-75-9 | NDMA | MutaMouse.CD2.lacZ80.HazfBR | 1perdayX28days | 28 |
| 16 | 5 | 62-75-9 | NDMA | MutaMouse.CD2.lacZ80.HazfBR | 1perdayX28days | 28 |
| 17 | 6 | 62-75-9 | NDMA | MutaMouse.CD2.lacZ80.HazfBR | 1perdayX28days | 28 |
| 18 | 1 | 62-75-9 | NDMA | MutaMouse.CD2.lacZ80.HazfBR | 1perdayX28days | 28 |
| 19 | 2 | 62-75-9 | NDMA | MutaMouse.CD2.lacZ80.HazfBR | 1perdayX28days | 28 |
| 20 | 3 | 62-75-9 | NDMA | MutaMouse.CD2.lacZ80.HazfBR | 1perdayX28days | 28 |
| 21 | 4 | 62-75-9 | NDMA | MutaMouse.CD2.lacZ80.HazfBR | 1perdayX28days | 28 |
| 22 | 5 | 62-75-9 | NDMA | MutaMouse.CD2.lacZ80.HazfBR | 1perdayX28days | 28 |
| 23 | 6 | 62-75-9 | NDMA | MutaMouse.CD2.lacZ80.HazfBR | 1perdayX28days | 28 |
| 24 | 1 | 62-75-9 | NDMA | MutaMouse.CD2.lacZ80.HazfBR | 1perdayX28days | 28 |
| 25 | 2 | 62-75-9 | NDMA | MutaMouse.CD2.lacZ80.HazfBR | 1perdayX28days | 28 |
| 26 | 3 | 62-75-9 | NDMA | MutaMouse.CD2.lacZ80.HazfBR | 1perdayX28days | 28 |
| 27 | 4 | 62-75-9 | NDMA | MutaMouse.CD2.lacZ80.HazfBR | 1perdayX28days | 28 |
| 28 | 5 | 62-75-9 | NDMA | MutaMouse.CD2.lacZ80.HazfBR | 1perdayX28days | 28 |
| 29 | 6 | 62-75-9 | NDMA | MutaMouse.CD2.lacZ80.HazfBR | 1perdayX28days | 28 |
| 30 | 1 | 62-75-9 | NDMA | MutaMouse.CD2.lacZ80.HazfBR | 1perdayX28days | 28 |
| 31 | 2 | 62-75-9 | NDMA | MutaMouse.CD2.lacZ80.HazfBR | 1perdayX28days | 28 |
| 32 | 3 | 62-75-9 | NDMA | MutaMouse.CD2.lacZ80.HazfBR | 1perdayX28days | 28 |
| 33 | 4 | 62-75-9 | NDMA | MutaMouse.CD2.lacZ80.HazfBR | 1perdayX28days | 28 |
| 34 | 5 | 62-75-9 | NDMA | MutaMouse.CD2.lacZ80.HazfBR | 1perdayX28days | 28 |
| 35 | 1 | 62-75-9 | NDMA | MutaMouse.CD2.lacZ80.HazfBR | 1perdayX28days | 28 |
| 36 | 2 | 62-75-9 | NDMA | MutaMouse.CD2.lacZ80.HazfBR | 1perdayX28days | 28 |
| 37 | 3 | 62-75-9 | NDMA | MutaMouse.CD2.lacZ80.HazfBR | 1perdayX28days | 28 |
| 38 | 4 | 62-75-9 | NDMA | MutaMouse.CD2.lacZ80.HazfBR | 1perdayX28days | 28 |
| 39 | 5 | 62-75-9 | NDMA | MutaMouse.CD2.lacZ80.HazfBR | 1perdayX28days | 28 |
| 40 | 1 | 62-75-9 | NDMA | MutaMouse.CD2.lacZ80.HazfBR | 1perdayX28days | 28 |
| 41 | 2 | 62-75-9 | NDMA | MutaMouse.CD2.lacZ80.HazfBR | 1perdayX28days | 28 |
| 42 | 3 | 62-75-9 | NDMA | MutaMouse.CD2.lacZ80.HazfBR | 1perdayX28days | 28 |
| 43 | 4 | 62-75-9 | NDMA | MutaMouse.CD2.lacZ80.HazfBR | 1perdayX28days | 28 |
| 44 | 5 | 62-75-9 | NDMA | MutaMouse.CD2.lacZ80.HazfBR | 1perdayX28days | 28 |
| 45 | 1 | 62-75-9 | NDMA | MutaMouse.CD2.lacZ80.HazfBR | 1perdayX1days | 1 |
| 46 | 2 | 62-75-9 | NDMA | MutaMouse.CD2.lacZ80.HazfBR | 1perdayX1days | 1 |
| 47 | 3 | 62-75-9 | NDMA | MutaMouse.CD2.lacZ80.HazfBR | 1perdayX1days | 1 |
| 48 | 4 | 62-75-9 | NDMA | MutaMouse.CD2.lacZ80.HazfBR | 1perdayX1days | 1 |
| 49 | 5 | 62-75-9 | NDMA | MutaMouse.CD2.lacZ80.HazfBR | 1perdayX1days | 1 |
| 50 | 1 | 62-75-9 | NDMA | MutaMouse.CD2.lacZ80.HazfBR | 1perdayX1days | 1 |
| 51 | 2 | 62-75-9 | NDMA | MutaMouse.CD2.lacZ80.HazfBR | 1perdayX1days | 1 |
| 52 | 3 | 62-75-9 | NDMA | MutaMouse.CD2.lacZ80.HazfBR | 1perdayX1days | 1 |
| 53 | 4 | 62-75-9 | NDMA | MutaMouse.CD2.lacZ80.HazfBR | 1perdayX1days | 1 |
| 54 | 1 | 759-73-9 | ENU | MutaMouse.CD2.lacZ80.HazfBR | NAN | NAN |
| 55 | 2 | 759-73-9 | ENU | MutaMouse.CD2.lacZ80.HazfBR | NAN | NAN |
| 56 | 3 | 759-73-9 | ENU | MutaMouse.CD2.lacZ80.HazfBR | NAN | NAN |

| row | sampling.time.h | dose | dose.units | tissue | MutationFrequency | type |
| --- | --- | --- | --- | --- | --- | --- |
| 1 | 72 | 0 | mg.kg.day | liver | 1.62E-07 | TwinStrandLacZPanel |
| 2 | 72 | 0 | mg.kg.day | liver | 1.20E-07 | TwinStrandLacZPanel |
| 3 | 72 | 0 | mg.kg.day | liver | 1.29E-07 | TwinStrandLacZPanel |
| 4 | 72 | 0 | mg.kg.day | liver | 1.53E-07 | TwinStrandLacZPanel |
| 5 | 72 | 0 | mg.kg.day | liver | 1.07E-07 | TwinStrandLacZPanel |
| 6 | 72 | 0 | mg.kg.day | liver | 1.58E-07 | TwinStrandLacZPanel |
| 7 | 72 | 0.02 | mg.kg.day | liver | 1.26E-07 | TwinStrandLacZPanel |
| 8 | 72 | 0.02 | mg.kg.day | liver | 1.38E-07 | TwinStrandLacZPanel |
| 9 | 72 | 0.02 | mg.kg.day | liver | 1.84E-07 | TwinStrandLacZPanel |
| 10 | 72 | 0.02 | mg.kg.day | liver | 1.50E-07 | TwinStrandLacZPanel |
| 11 | 72 | 0.02 | mg.kg.day | liver | 1.31E-07 | TwinStrandLacZPanel |
| 12 | 72 | 0.07 | mg.kg.day | liver | 1.27E-07 | TwinStrandLacZPanel |
| 13 | 72 | 0.07 | mg.kg.day | liver | 1.86E-07 | TwinStrandLacZPanel |
| 14 | 72 | 0.07 | mg.kg.day | liver | 1.41E-07 | TwinStrandLacZPanel |
| 15 | 72 | 0.07 | mg.kg.day | liver | 1.62E-07 | TwinStrandLacZPanel |
| 16 | 72 | 0.07 | mg.kg.day | liver | 1.26E-07 | TwinStrandLacZPanel |
| 17 | 72 | 0.07 | mg.kg.day | liver | 1.45E-07 | TwinStrandLacZPanel |
| 18 | 72 | 0.19 | mg.kg.day | liver | 2.07E-07 | TwinStrandLacZPanel |
| 19 | 72 | 0.19 | mg.kg.day | liver | 2.25E-07 | TwinStrandLacZPanel |
| 20 | 72 | 0.19 | mg.kg.day | liver | 1.58E-07 | TwinStrandLacZPanel |
| 21 | 72 | 0.19 | mg.kg.day | liver | 1.76E-07 | TwinStrandLacZPanel |
| 22 | 72 | 0.19 | mg.kg.day | liver | 2.45E-07 | TwinStrandLacZPanel |
| 23 | 72 | 0.19 | mg.kg.day | liver | 1.95E-07 | TwinStrandLacZPanel |
| 24 | 72 | 0.36 | mg.kg.day | liver | 4.58E-07 | TwinStrandLacZPanel |
| 25 | 72 | 0.36 | mg.kg.day | liver | 3.35E-07 | TwinStrandLacZPanel |
| 26 | 72 | 0.36 | mg.kg.day | liver | 4.95E-07 | TwinStrandLacZPanel |
| 27 | 72 | 0.36 | mg.kg.day | liver | 2.96E-07 | TwinStrandLacZPanel |
| 28 | 72 | 0.36 | mg.kg.day | liver | 3.88E-07 | TwinStrandLacZPanel |
| 29 | 72 | 0.36 | mg.kg.day | liver | 3.97E-07 | TwinStrandLacZPanel |
| 30 | 72 | 1.1 | mg.kg.day | liver | 5.93E-07 | TwinStrandLacZPanel |
| 31 | 72 | 1.1 | mg.kg.day | liver | 6.55E-07 | TwinStrandLacZPanel |
| 32 | 72 | 1.1 | mg.kg.day | liver | 6.94E-07 | TwinStrandLacZPanel |
| 33 | 72 | 1.1 | mg.kg.day | liver | 7.13E-07 | TwinStrandLacZPanel |
| 34 | 72 | 1.1 | mg.kg.day | liver | 7.70E-07 | TwinStrandLacZPanel |
| 35 | 72 | 2 | mg.kg.day | liver | 6.74E-07 | TwinStrandLacZPanel |
| 36 | 72 | 2 | mg.kg.day | liver | 7.34E-07 | TwinStrandLacZPanel |
| 37 | 72 | 2 | mg.kg.day | liver | 7.79E-07 | TwinStrandLacZPanel |
| 38 | 72 | 2 | mg.kg.day | liver | 6.32E-07 | TwinStrandLacZPanel |
| 39 | 72 | 2 | mg.kg.day | liver | 7.68E-07 | TwinStrandLacZPanel |
| 40 | 72 | 4 | mg.kg.day | liver | 8.87E-07 | TwinStrandLacZPanel |
| 41 | 72 | 4 | mg.kg.day | liver | 7.85E-07 | TwinStrandLacZPanel |
| 42 | 72 | 4 | mg.kg.day | liver | 7.92E-07 | TwinStrandLacZPanel |
| 43 | 72 | 4 | mg.kg.day | liver | 6.73E-07 | TwinStrandLacZPanel |
| 44 | 72 | 4 | mg.kg.day | liver | 7.43E-07 | TwinStrandLacZPanel |
| 45 | 720 | 5 | mg.kg | liver | 2.01E-07 | TwinStrandLacZPanel |
| 46 | 720 | 5 | mg.kg | liver | 2.07E-07 | TwinStrandLacZPanel |
| 47 | 720 | 5 | mg.kg | liver | 2.11E-07 | TwinStrandLacZPanel |
| 48 | 720 | 5 | mg.kg | liver | 2.10E-07 | TwinStrandLacZPanel |
| 49 | 720 | 5 | mg.kg | liver | 2.66E-07 | TwinStrandLacZPanel |
| 50 | 720 | 10 | mg.kg | liver | 2.80E-07 | TwinStrandLacZPanel |
| 51 | 720 | 10 | mg.kg | liver | 2.87E-07 | TwinStrandLacZPanel |
| 52 | 720 | 10 | mg.kg | liver | 2.80E-07 | TwinStrandLacZPanel |
| 53 | 720 | 10 | mg.kg | liver | 2.95E-07 | TwinStrandLacZPanel |
| 54 | NAN | 50 | mg.kg | liver | 5.30E-07 | TwinStrandLacZPanel |
| 55 | NAN | 50 | mg.kg | liver | 4.33E-07 | TwinStrandLacZPanel |
| 56 | NAN | 50 | mg.kg | liver | 3.65E-07 | TwinStrandLacZPanel |

| row | sampling.time.h | dose | dose.units | tissue | MutationFrequency | type |
| --- | --- | --- | --- | --- | --- | --- |
| 1 | 72 | 0 | mg.kg.day | liver | 5.31E-08 | TwinStrandMouseMutagenesisPanel |
| 2 | 72 | 0 | mg.kg.day | liver | 6.62E-08 | TwinStrandMouseMutagenesisPanel |
| 3 | 72 | 0 | mg.kg.day | liver | 5.35E-08 | TwinStrandMouseMutagenesisPanel |
| 4 | 72 | 0 | mg.kg.day | liver | 4.25E-08 | TwinStrandMouseMutagenesisPanel |
| 5 | 72 | 0 | mg.kg.day | liver | 4.74E-08 | TwinStrandMouseMutagenesisPanel |
| 6 | 72 | 0 | mg.kg.day | liver | 3.81E-08 | TwinStrandMouseMutagenesisPanel |
| 7 | 72 | 0.02 | mg.kg.day | liver | 4.30E-08 | TwinStrandMouseMutagenesisPanel |
| 8 | 72 | 0.02 | mg.kg.day | liver | 3.62E-08 | TwinStrandMouseMutagenesisPanel |
| 9 | 72 | 0.02 | mg.kg.day | liver | 4.19E-08 | TwinStrandMouseMutagenesisPanel |
| 10 | 72 | 0.02 | mg.kg.day | liver | 5.04E-08 | TwinStrandMouseMutagenesisPanel |
| 11 | 72 | 0.02 | mg.kg.day | liver | 4.61E-08 | TwinStrandMouseMutagenesisPanel |
| 12 | 72 | 0.07 | mg.kg.day | liver | 5.54E-08 | TwinStrandMouseMutagenesisPanel |
| 13 | 72 | 0.07 | mg.kg.day | liver | 7.27E-08 | TwinStrandMouseMutagenesisPanel |
| 14 | 72 | 0.07 | mg.kg.day | liver | 6.43E-08 | TwinStrandMouseMutagenesisPanel |
| 15 | 72 | 0.07 | mg.kg.day | liver | 7.14E-08 | TwinStrandMouseMutagenesisPanel |
| 16 | 72 | 0.07 | mg.kg.day | liver | 3.67E-08 | TwinStrandMouseMutagenesisPanel |
| 17 | 72 | 0.07 | mg.kg.day | liver | 5.48E-08 | TwinStrandMouseMutagenesisPanel |
| 18 | 72 | 0.19 | mg.kg.day | liver | 1.15E-07 | TwinStrandMouseMutagenesisPanel |
| 19 | 72 | 0.19 | mg.kg.day | liver | 1.08E-07 | TwinStrandMouseMutagenesisPanel |
| 20 | 72 | 0.19 | mg.kg.day | liver | 7.00E-08 | TwinStrandMouseMutagenesisPanel |
| 21 | 72 | 0.19 | mg.kg.day | liver | 7.50E-08 | TwinStrandMouseMutagenesisPanel |
| 22 | 72 | 0.19 | mg.kg.day | liver | 1.34E-07 | TwinStrandMouseMutagenesisPanel |
| 23 | 72 | 0.19 | mg.kg.day | liver | 9.93E-08 | TwinStrandMouseMutagenesisPanel |
| 24 | 72 | 0.36 | mg.kg.day | liver | 3.34E-07 | TwinStrandMouseMutagenesisPanel |
| 25 | 72 | 0.36 | mg.kg.day | liver | 1.96E-07 | TwinStrandMouseMutagenesisPanel |
| 26 | 72 | 0.36 | mg.kg.day | liver | 3.40E-07 | TwinStrandMouseMutagenesisPanel |
| 27 | 72 | 0.36 | mg.kg.day | liver | 2.08E-07 | TwinStrandMouseMutagenesisPanel |
| 28 | 72 | 0.36 | mg.kg.day | liver | 3.11E-07 | TwinStrandMouseMutagenesisPanel |
| 29 | 72 | 0.36 | mg.kg.day | liver | 2.65E-07 | TwinStrandMouseMutagenesisPanel |
| 30 | 72 | 1.1 | mg.kg.day | liver | 5.00E-07 | TwinStrandMouseMutagenesisPanel |
| 31 | 72 | 1.1 | mg.kg.day | liver | 6.11E-07 | TwinStrandMouseMutagenesisPanel |
| 32 | 72 | 1.1 | mg.kg.day | liver | 6.40E-07 | TwinStrandMouseMutagenesisPanel |
| 33 | 72 | 1.1 | mg.kg.day | liver | 6.64E-07 | TwinStrandMouseMutagenesisPanel |
| 34 | 72 | 1.1 | mg.kg.day | liver | 7.13E-07 | TwinStrandMouseMutagenesisPanel |
| 35 | 72 | 2 | mg.kg.day | liver | 6.45E-07 | TwinStrandMouseMutagenesisPanel |
| 36 | 72 | 2 | mg.kg.day | liver | 6.46E-07 | TwinStrandMouseMutagenesisPanel |
| 37 | 72 | 2 | mg.kg.day | liver | 6.44E-07 | TwinStrandMouseMutagenesisPanel |
| 38 | 72 | 2 | mg.kg.day | liver | 5.98E-07 | TwinStrandMouseMutagenesisPanel |
| 39 | 72 | 2 | mg.kg.day | liver | 7.63E-07 | TwinStrandMouseMutagenesisPanel |
| 40 | 72 | 4 | mg.kg.day | liver | 8.90E-07 | TwinStrandMouseMutagenesisPanel |
| 41 | 72 | 4 | mg.kg.day | liver | 5.02E-07 | TwinStrandMouseMutagenesisPanel |
| 42 | 72 | 4 | mg.kg.day | liver | 7.83E-07 | TwinStrandMouseMutagenesisPanel |
| 43 | 72 | 4 | mg.kg.day | liver | 6.10E-07 | TwinStrandMouseMutagenesisPanel |
| 44 | 72 | 4 | mg.kg.day | liver | 5.53E-07 | TwinStrandMouseMutagenesisPanel |
| 45 | 720 | 5 | mg.kg | liver | 1.11E-07 | TwinStrandMouseMutagenesisPanel |
| 46 | 720 | 5 | mg.kg | liver | 1.39E-07 | TwinStrandMouseMutagenesisPanel |
| 47 | 720 | 5 | mg.kg | liver | 1.23E-07 | TwinStrandMouseMutagenesisPanel |
| 48 | 720 | 5 | mg.kg | liver | 1.46E-07 | TwinStrandMouseMutagenesisPanel |
| 49 | 720 | 5 | mg.kg | liver | 1.88E-07 | TwinStrandMouseMutagenesisPanel |
| 50 | 720 | 10 | mg.kg | liver | 2.65E-07 | TwinStrandMouseMutagenesisPanel |
| 51 | 720 | 10 | mg.kg | liver | 2.15E-07 | TwinStrandMouseMutagenesisPanel |
| 52 | 720 | 10 | mg.kg | liver | 2.59E-07 | TwinStrandMouseMutagenesisPanel |
| 53 | 720 | 10 | mg.kg | liver | 2.65E-07 | TwinStrandMouseMutagenesisPanel |
| 54 | NAN | 50 | mg.kg | liver | 5.66E-07 | TwinStrandMouseMutagenesisPanel |
| 55 | NAN | 50 | mg.kg | liver | 4.17E-07 | TwinStrandMouseMutagenesisPanel |
| 56 | NAN | 50 | mg.kg | liver | 4.00E-07 | TwinStrandMouseMutagenesisPanel |

